# A trimeric Rab7 GEF controls NPC1-dependent lysosomal cholesterol export

**DOI:** 10.1101/835686

**Authors:** Dick J.H. van den Boomen, Agata Sienkiewicz, Ilana Berlin, Marlieke L.M. Jongsma, Daphne M. van Elsland, J. Paul Luzio, Jacques J.C. Neefjes, Paul J. Lehner

## Abstract

Cholesterol import in mammalian cells is mediated by the LDL receptor pathway. Here, using an endogenous cholesterol reporter in a genome-wide CRISPR screen we identify >70 genes involved in LDL-cholesterol import. We characterise C18orf8 as a core component of the mammalian Mon1-Ccz1 guanidine exchange factor (GEF) for Rab7, required for complex stability and function. *C18orf8*-deficient cells lack Rab7 activation and show severe defects in late endosome morphology and endosomal LDL trafficking, resulting in cellular cholesterol deficiency. Unexpectedly, free cholesterol accumulates within swollen lysosomes, suggesting a critical additional defect in lysosomal cholesterol export. We find that active Rab7 interacts with the NPC1 cholesterol transporter and licenses lysosomal cholesterol export. This process is abolished in *C18orf8*-, *Ccz1*- and *Mon1A/B*-deficient cells and restored by a constitutively active Rab7. The trimeric Mon1-Ccz1-C18orf8 (MCC) GEF therefore plays a central role in cellular cholesterol homeostasis coordinating Rab7 activation, endosomal LDL trafficking and NPC1-dependent lysosomal cholesterol export.

## Introduction

Cholesterol is an essential component of all mammalian lipid membranes and disturbances in its homeostasis associate with a variety of diseases, including Niemann Pick type C, familial hypercholesterolemia, atherosclerosis and obesity. Cellular cholesterol homeostasis is transcriptionally regulated by the SREBP2 (sterol response element binding protein 2) transcription factor. When cholesterol is abundant SREBP2 and its trafficking factor SCAP are retained in the endoplasmic reticulum (ER), whereas upon sterol depletion they are released to the Golgi, where SREBP2 is cleaved and traffics to the nucleus to activate the transcription of genes involved in *de novo* cholesterol biosynthesis (e.g. *HMGCS1*, *HMGCR*, *SQLE*) and LDL-cholesterol uptake (e.g. *LDLR*, *NPC1/2*) (1). SREBP2 thus couples cholesterol import and biosynthesis to cellular cholesterol availability.

Cholesterol is taken up into mammalian cells in the form of low-density lipoprotein (LDL) particles, composed predominantly of cholesterol esters (CEs), cholesterol and apolipoprotein B-100 (2). LDL binds to the cell surface LDL receptor (LDLR) and the LDLR-LDL complex is internalised via clathrin-mediated endocytosis (3). In the acidic pH of sorting endosomes, LDL dissociates from LDLR, which recycles back to the plasma membrane. LDL is itself transported to a late endosomal / lysosomal (LE/Ly) compartment, where cholesterol is released from CEs by lysosomal acid lipase (LAL/LIPA). Free cholesterol then binds the Niemann Pick C2 (NPC2) carrier protein and is transferred to the Niemann Pick C1 (NPC1) transporter for subsequent export to other organelles (4).

NPC1 is a central mediator of lysosomal cholesterol export. Mutations in *NPC1* and *NPC2* cause Niemann Pick type C, a lethal lysosomal storage disease characterised by lysosomal cholesterol accumulation in multiple organ systems. *NPC1* encodes a complex polytopic glycoprotein embedded in the lysosomal membrane (5). Cholesterol-loaded NPC2 binds the NPC1 middle luminal domain (6, 7) and delivers its cargo to the mobile N-terminal domain (8, 9), from where cholesterol is likely transferred to the sterol-sensing domain (SSD). Whether cholesterol is transported through NPC1 in a channel-like manner or simply inserted into the luminal membrane leaflet remains unclear (10). NPC1 is believed to be constitutively active and no regulators of its activity have been identified. Once transported across the lysosomal membrane, cholesterol transfer to other organelles is mediated by lipid transfer proteins (LTPs) at inter-organelle membrane contact sites (MCS) (11). Although a variety of proteins have been implicated in MCS formation with LE/Ly (12–15), the identity of the direct carrier transporting cholesterol from LE/Ly remains controversial.

The LE/Ly compartment plays a central role in the cellular LDL-cholesterol uptake pathway. LE homeostasis and substrate trafficking are regulated by the small GTPase Rab7 (16). Rab7 activity is controlled by its nucleotide status: its activation requires GDP-to-GTP exchange by a guanidine exchange factor (GEF), whereas GTP hydrolysis induced by GTPase activating proteins (GAPs) triggers Rab7 inactivation. Active GTP-bound Rab7 associates with LE membranes and recruits Rab7 effector proteins which in mammalian cells include the endosome motility factor RILP (17, 18), the cholesterol binding protein ORP1L (19, 20), the VPS34 phosphatidylinositol (PtdIns) 3-kinase regulators Rubicon and WDR91 (21, 22) and the retromer components VPS26 and VPS35 (23, 24).

Temporal and spatial control of Rab7 activation is of major importance. At least three GAPs – TBC1D5, TBC1D15 and Armus (24–26) are implicated in Rab7 inactivation, whereas activation of the yeast Rab7 homologue Ypt7 is mediated by the Mon1-Ccz1 (MC1) complex (27, 28). MC1 is recruited to endosomal membranes by the phospholipid PtdIns(3)P (29) and its activation of Rab7 drives Rab5-to-Rab7 conversion, endosome maturation and fusion with the vacuolar/lysosomal compartment (30–33). A partial crystal structure of the *C. thermophilum* MC1-Ypt7 complex suggests a GEF model in which MC1 binding to Ypt7 induces magnesium expulsion and GDP dissociation from the Ypt7 active site (34). Mammals have two Mon1 orthologues – Mon1A and Mon1B – and a single Ccz1 orthologue. Although mammalian MC1 also likely acts as a Rab7 GEF (35, 36), its *in vivo* function remains poorly characterised.

In this study, we performed a genome-wide CRISPR screen for essential genes in cholesterol homeostasis using an endogenous SREBP2-dependent cholesterol reporter. We identified >70 genes mainly involved in the LDL-cholesterol uptake pathway, including the poorly characterised *C18orf8*. C18orf8 is a novel core component of the mammalian MC1 complex, essential for complex stability and function. *C18orf8*-deficient cells exhibit severe defects in Rab7 activation and LDL trafficking, concomitant with swelling of the LE/Ly compartment and marked lysosomal cholesterol accumulation. We show that active Rab7 interacts with the NPC1 cholesterol transporter to license lysosomal cholesterol export - a pathway deficient in *C18orf8-*, *Ccz1*- and *Mon1A/B*-deficient cells and restored by a constitutively active Rab7 (Q67L). Our findings therefore identify a central role for the trimeric <u>M</u>on1-<u>C</u>cz1-<u>C</u>18orf8 (MCC) GEF in cellular LDL-cholesterol uptake, coordinating Rab7 activation with LDL trafficking and NPC1-dependent lysosomal cholesterol export.

## Results

### Generation of an endogenous, SREBP2-dependent, fluorescent cholesterol reporter

To screen for genes that maintain cellular cholesterol homeostasis, we engineered a cell line expressing an endogenous fluorescent reporter responsive to intracellular cholesterol levels. HMG-CoA synthase 1 (HMGCS1) catalyses the second step of cholesterol biosynthesis, the conversion of acetoacetyl-CoA and acetyl-CoA to HMG-CoA. With two sterol-response elements (SRE) in its promoter, HMGCS1 expression is highly SREBP2 responsive and thus cholesterol sensitive (37), rendering it well suited to monitor cellular cholesterol levels. We used CRISPR technology to knock-in the bright fluorescent protein Clover (38) into the endogenous *HMGCS1* locus yielding an endogenous HMGCS1-Clover fusion protein (Fig 1A, S1A). Sterol depletion of the *HMGCS1-Clover* cell line, using mevastatin and lipoprotein-depleted serum (LPDS), increased basal HMGCS1-Clover expression 9-fold (Fig 1C, S1B, S1C) and revealed the expected cytoplasmic localisation of the HMGCS1-Clover fusion protein (Fig 1B). Expression returned to baseline following overnight sterol repletion (Fig S1D). Sterol-depletion induced HMGCS1-Clover expression was abolished upon knockout of *SREBP2*, but not its close relative *SREBP1* (Fig 1C, S1E). *SREBP2*-deficiency also slightly decreased steady-state HMGCS1-Clover levels (Fig S1F), suggesting low-level SREBP2 activation under standard growth conditions. Unlike HMG-CoA reductase (HMGCR), HMGCS1 expression is not regulated by sterol-induced degradation (Fig S1G). *HMGCS1-Clover* is therefore a sensitive endogenous, SREBP2-dependent, transcriptional reporter for intracellular cholesterol levels.

**Figure 1:**
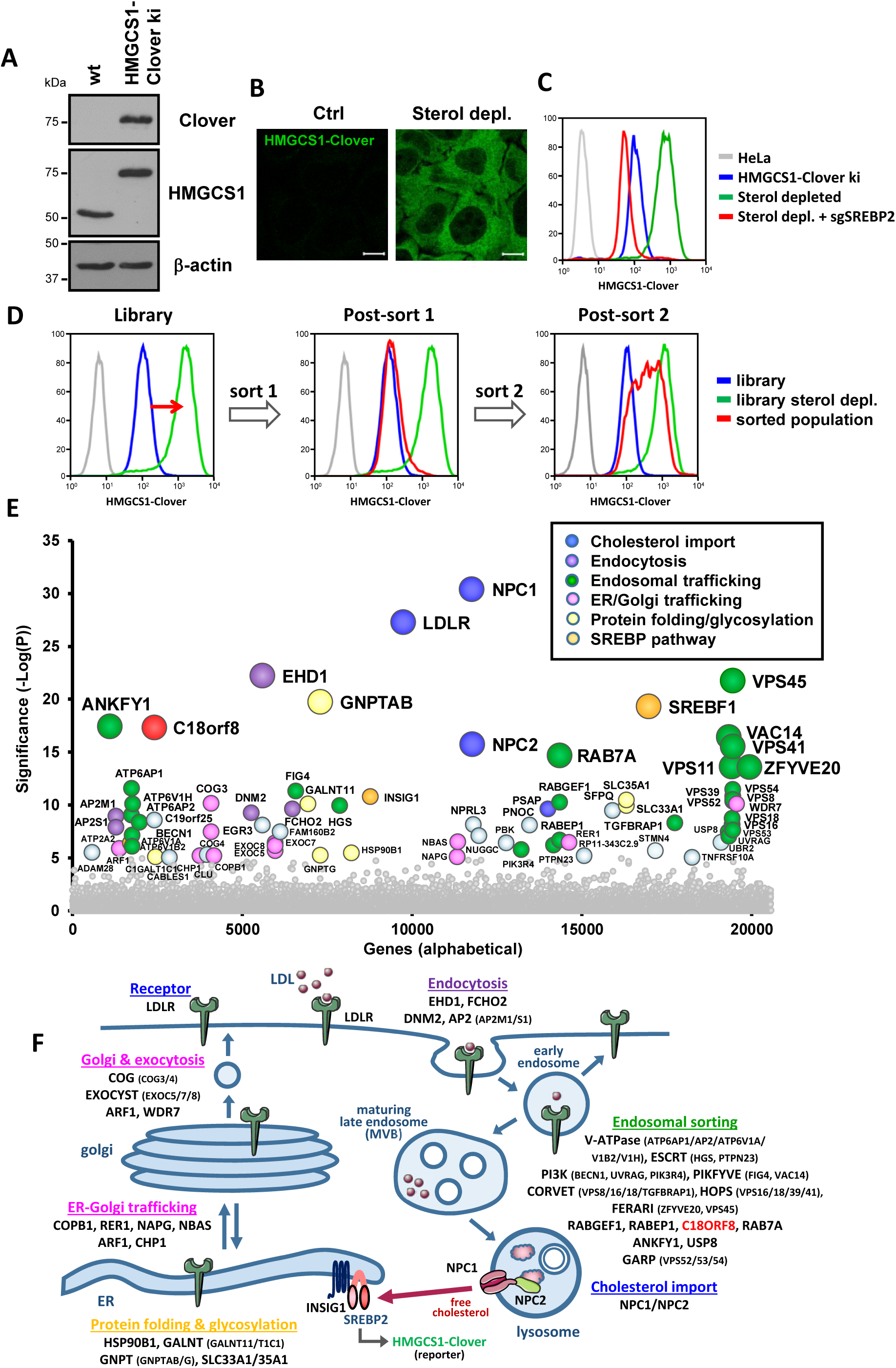
A genome-wide CRISPR screen identifies essential factors in cellular cholesterol homeostasis. (**A** - **C**) Characterisation of a HMGCS1-Clover CRISPR knock-in in HeLa cells. (**A**) Immunoblotting of a reporter clone shows the endogenous HMGCS1-Clover fusion protein is detected by both HMGCS1- and GFP-specific antibody staining. (**B**) Sterol depletion induces HMGCS1-Clover expression. HeLa HMGCS1-Clover cells were sterol depleted overnight and HMGCS1-Clover expression was analysed by confocal microscopy. (**C**) HMGCS1 upregulation is SREBP2-dependent. HeLa HMGCS1-Clover CAS9 cells were transfected with sgRNA against *SREBF2*, and after 7 days, their response to sterol depletion was determined by flow cytometry. (**D** - **F**) A genome-wide CRISPR screen identifies essential factors in cellular cholesterol homeostasis. (**D**) HeLa HMGCS1-Clover/Cas9 cells were transduced with a genome-wide sgRNA library (22,000 sgRNAs). Rare HMGCS1-Clover^high^ cells were isolated using two rounds of cell sorting, resulting in a 60% enriched HMGCS1-Clover^high^ population. (**E**) Illumina sequencing of sgRNAs in the isolated HMGCS1-Clover^high^ population shows sgRNA enrichment for genes involved in cholesterol uptake (LDLR, NPC1, NPC2; blue), protein folding and glycosylation (yellow), membrane trafficking of LDLR/LDL (pink, purple, green) and SREBP2 function (orange). Genes with sgRNA enrichment at P<10^-5^ are indicated. (**F**) A schematic representation of hits from the genome-wide CRISPR screen for cholesterol regulators highlights the central role for LDL-cholesterol import for cellular cholesterol homeostasis. Hits involved in pathways other than those indicated can be found in **Table S1**.

### A genome-wide CRISPR screen identifies components essential for cellular cholesterol homeostasis

To identify genes critical to the maintenance of cellular cholesterol levels, we undertook a genome-wide CRISPR screen using our HMGCS1-Clover reporter as an intracellular cholesterol sensor. CRISPR knockout of these genes is predicted to induce spontaneous cholesterol depletion, SREBP2 activation and HMGCS1-Clover upregulation, despite sterol replete (high cholesterol) culture conditions. These cells thus develop a spontaneous HMGCS1-Clover^high^ phenotype. *HMGCS1-Clover* reporter cells were targeted with a genome-wide CRISPR library containing 10 sgRNAs per gene (total sgRNA size 220,000) (39) and were sorted in two successive rounds of flow cytometry for the rare HMGCS1-Clover^high^ phenotype (Fig 1D). This yielded an enriched population with a 60% HMGCS1-Clover^high^ phenotype under sterol replete conditions. Next-generation sequencing of the population revealed sgRNA enrichment for 76 genes (P<10^-5^) (Fig 1E, **Table S1**). The results of our HMGCS1-Clover screen emphasise the central role of LDL-cholesterol uptake and membrane trafficking in cellular cholesterol homeostasis, with top hits including *LDLR* and *NPC1/2*.

Functionally, the hits can be grouped in categories (Fig 1E) that include: **i)** protein folding and glycosylation (**yellow**, e.g. *HSP90B1*, *GNTAP*, *SLC35A1*); **ii)** trafficking through the early secretory pathway (**pink**, e.g. *ARF1*, *CHP1*, *COG3/4*, *EXOC5/7/9*); **iii)** endocytosis (**purple**, e.g. *AP2M1/S1*, *EHD1*, *FCHO2*, *DNM2*); **iv)** endosomal trafficking (**green,** e.g. *RabGEF1*, *Rab7A*, V-ATPase (*ATP6AP1/AP2V1A/V1B2/V1H*), ESCRT (*PTPN23, HRS*), CORVET (*VPS8/16/18/TGFBRAP1*), HOPS (*VPS16/18/39/41*), FERARI (*ZFYVE20/RBSN, VPS45),* PI3K (*BECN, UVRAG, PIK3R4*), PIKfyve (*FIG4, VAC14*)); and **v)** cholesterol import (**blue**, e.g. *NPC1/2*). These categories reflect successive stages in LDL-cholesterol import (Fig 1F), respectively: **i)** LDLR folding and glycosylation; **ii)** trafficking of LDLR to the cell surface; **iii)** endocytosis of the LDL-LDLR complex; **iv)** trafficking of internalised LDL to the LE/Ly compartment; and **v)** lysosomal cholesterol release and NPC1-dependent lysosomal cholesterol export. Besides the LDLR pathway, our screen identified components of the SREBP transcriptional machinery (Fig 1E, **orange**, *INSIG1, SREBP1*) and several poorly characterised gene products (**grey/red** e.g. *CLU*/*APOJ*, *EGR3*, *ADAM28, FAM160B2*, *STMN4, C18orf8*). To validate our screen results, we confirmed a subset of hits (*LDLR, NPC1, AP2μ1, ANKFY1, VPS16, RBSN/ZFYVE20, INSIG1, SREBF1*) using individual sgRNAs. All sgRNA-treated cells showed elevated HMGCS1-Clover expression (Fig S2), suggesting defective LDL-cholesterol import and/or spontaneous SREBP2 activation.

**Figure 2:**
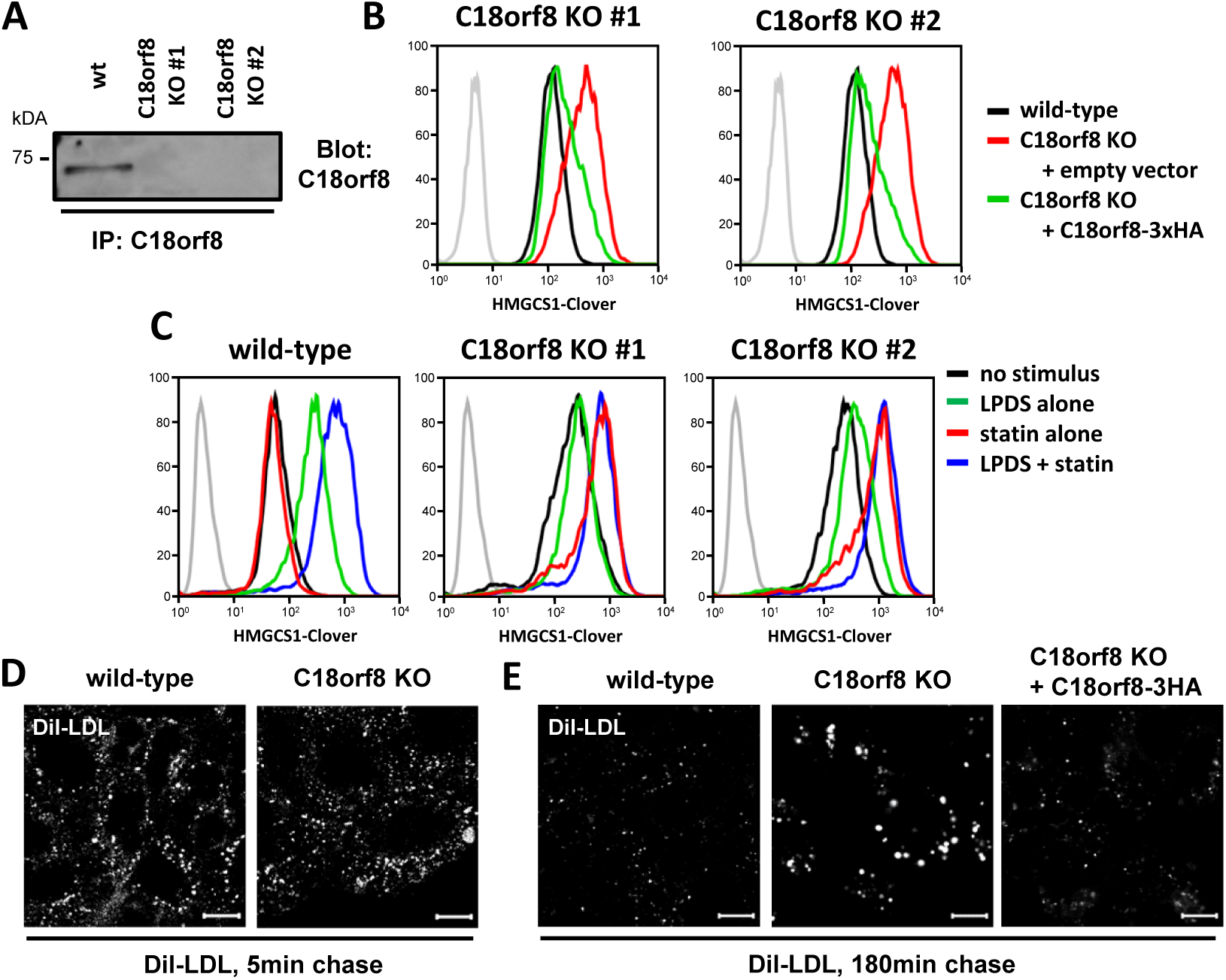
C18orf8 is required for endosomal LDL-cholesterol uptake. (**A**, **B**) C18orf8-deficient cells show spontaneous cholesterol deficiency. (**A**) Wild-type and *C18orf8*-deficient clones were lysed, endogenous C18orf8 immunoprecipitated, and detected by a C18orf8-specific antibody. (**B**) *C18orf8*-deficient HMGCS1-Clover clones were transduced with HA-tagged *C18orf8* (C18orf8-3xHA; green lines) or empty vector (red lines) and HMGCS1-Clover expression was determined by flow cytometry at day 18 using wild-type HMGCS1-Clover cells as a control (black lines). (**C**) *C18orf8*-deficient cells are dependent on endogenous cholesterol biosynthesis. Wild-type and *C18orf8*-deficient HMGCS1-Clover cells were either LPDS depleted to block exogenous LDL-cholesterol uptake (green lines), treated with mevastatin to block endogenous cholesterol biosynthesis (red lines), or a combination of both treatments (blue lines), after which cells were analysed by flow cytometry for HMGCS1-Clover expression. (**E, F**) *C18orf8*-deficient cells show defective endo-lysosomal LDL degradation. Wild-type, *C18orf8*-deficient or *C18orf8*-deficient cells complemented with C18orf8-3xHA were starved for 1 hour and pulse-labelled with fluorescent Dil-LDL, grown for the 5 min (**E**) or 180 min (**F**), fixed and visualised by confocal microscopy. Exposure times were kept constant between individual conditions at given time points. Scale bars = 10µm.

Our cholesterol reporter screen therefore provides a comprehensive overview and unique insight into known and unknown factors essential for LDL-cholesterol import into mammalian cells (Fig 1F), and emphasises the critical role of endo-/lysosomal trafficking within this process.

### *C18orf8* is required for endo-lysosomal LDL-cholesterol uptake

A prominent uncharacterised gene in our screen is *C18orf8* (Fig 1E, **red**), which encodes a 72kDa soluble protein, conserved from animals to plants with no identifiable yeast orthologue. Structure prediction (Phyre2, HHPred) suggests high structural similarity to the N-terminus of clathrin heavy chain, with an N-terminal β-propeller composed of WD40 repeats and a C-terminal α-helical domain (Fig S4E). Organelle proteomics have previously identified C18orf8 as a lysosome (40) and endosome (41) associated protein.

To further characterise *C18orf8* function, we generated *C18orf8* knockout clones using two independent sgRNAs (Fig 2A). Consistent with our screening results, *C18orf8-*deficient cell clones showed elevated HMGCS1-Clover expression under sterol-replete culture conditions (Fig 2B, **red line**) – implying spontaneous cholesterol depletion – and this phenotype was reversed upon re-expression of HA-tagged *C18orf8* (C18orf8-3xHA, **green line**). Cholesterol deficiency could result from defective exogenous LDL-cholesterol uptake or defective endogenous biosynthesis. To differentiate these pathways, we inhibited exogenous uptake by culturing cells in LPDS or blocked endogenous biosynthesis with mevastatin (Fig 2C). Wild-type reporter HeLa cells showed no response to mevastatin alone (Fig 2C, **left panel**, **red line**), upregulated HMGCS1-Clover in LPDS (**green line**) and showed maximum reporter induction in LPDS and mevastatin (**blue line**). In contrast, *C18orf8*-deficient cells showed elevated steady-state HMGCS1-Clover expression (Fig 2C, **middle/right panels, black lines**), were unresponsive to LPDS (**green lines**), but addition of mevastatin alone induced maximum HMGCS1-Clover induction (**red lines**) with no further increase in LPDS + mevastatin (**blue line**). These results show that while wild-type HeLa cells rely predominantly on external cholesterol and switch to endogenous biosynthesis when LDL is unavailable, *C18orf8*-deficient cells are completely reliant on endogenous cholesterol biosynthesis under all conditions. This effect is likely explained by a defect in LDL-cholesterol uptake.

*C18orf8*-deficient cells showed normal LDLR cell surface expression (Fig S3A) and uptake of fluorescent Dil-LDL (Fig 2D). However, whereas in wild-type cells, Dil-LDL fluorescence disappeared within 3 hours of pulse-labelling, in *C18orf8*-deficient cells Dil-LDL accumulated during the 3-hour chase (Fig 2E), a phenotype rescued upon complementation with C18orf8-3xHA. To determine whether this defect is LDL-specific, or derives from a general endolysosomal trafficking defect, we stimulated cells with fluorescent-labelled EGF. Similar to Dil-LDL, EGF was degraded within 3 hours in wild-type cells, but accumulated during the 3h chase in *C18orf8*-deficient cells (**Fig S3B**). *C18orf8*-deficient cells thus show a general defect in endo-lysosomal degradation that affects both LDL and EGF and results in their reliance on *de novo* cholesterol biosynthesis for cholesterol supply.

### *C18orf8-*deficient cells are defective in late endosome morphology and early-to-late substrate trafficking

Consistent with a defect in LDL/EGF degradation, *C18orf8*-deficient cells showed a severe disruption of endosome morphology, in particular of the LE/Ly compartment (Fig 3A, 3B, S3C). Perinuclear Rab7+ LEs were markedly swollen and surrounded by equally swollen LAMP1+ LE/Lys, with EEA1+/Rab5+ EEs clustered between Rab7+ LEs in the perinuclear region. By electron microscopy (EM), the swollen LE/Lys appeared as enlarged multivesicular bodies (MVBs) with a high intraluminal vesicle (ILV) content and a 3-fold increase in average diameter (Fig S3D). Complementation of *C18orf8*-deficient cells with C18orf8xHA was slow, but after 14 days restored endosome morphology (Fig S3E) and function (Fig 2E) back to wild-type. This prolonged recovery likely reflects the severity of the cellular phenotype.

**Figure 3:**
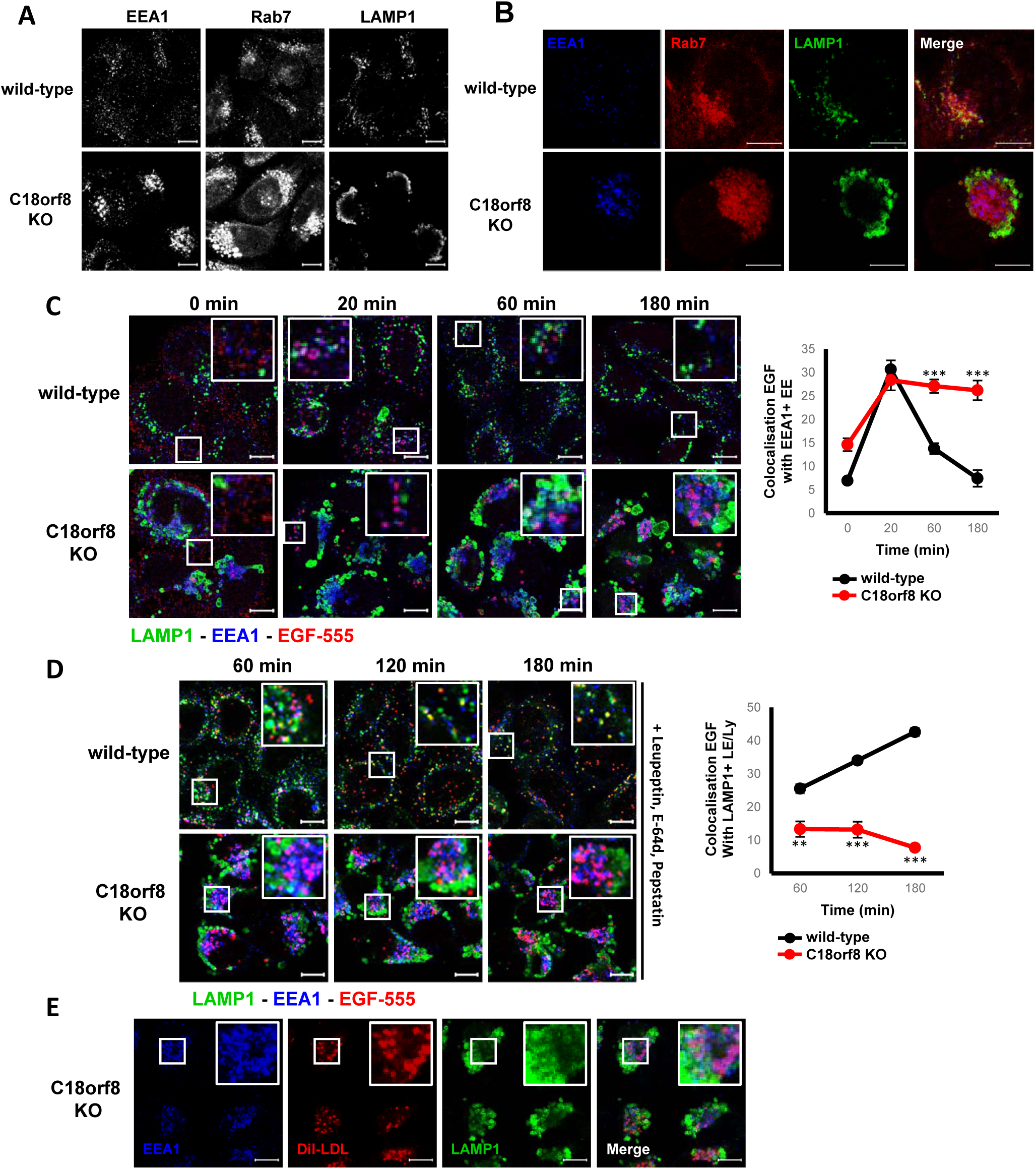
*C18orf8*-deficient cells show severe defects in late endosome morphology and early-to-late endosomal trafficking. (**A**, **B**) *C18orf8*-deficient cells show clustering of EE and swelling of LE/Ly. Confocal microscopy comparing EEA1, Rab7 and LAMP1in wild-type versus *C18orf8*-deficient cells. (**C - E**) *C18orf8*-deficient cells have severe defects in early-to-late endosomal trafficking. (**C, D**) Wild-type and *C18orf8*-deficient cells were starved for 1 hour, pulse-labelled with AlexaFluor555-conjugated EGF, incubated for the indicated times, fixed and stained for EEA1 (blue) and LAMP1 (green). In (**D**) a cocktail of protease inhibitors (Leupeptin/E-64d/Leupeptin) was added during starve and chase to block EGF degradation. EEA1+ and LAMP1+ vesicles were identified using Volocity software and the percentage of EGF co-localising with these structures was determined from 5-6 images per condition with at least eight cells per image (** P<0.01, *** P<0.001). (**E**) *C18orf8*-deficient cells were stimulated with Dil-LDL, chased for 3 hours, fixed and stained for EEA1 (blue) and LAMP1 (green). Scale bars = 10µm.

To assess how this abnormal morphology affects substrate trafficking, we pulse-labelled wild-type and *C18orf8*-deficient cells with fluorescent-labelled EGF. In both cell lines EGF moved to EEA1+ EE within ∼20min (Fig 3C). However, whereas in wild-type cells EGF disappeared from EE at 1-3 hours and was degraded (Fig 3C, **top panel**), in *C18orf8*-deficient cells no degradation occurred and EGF remained in EEA1+ EE for the duration of the 3 hours chase (**bottom panel**), implying a defect in early-to-late substrate trafficking. Indeed, inhibiting EGF degradation with Leupeptin, E-64d and Pepstatin showed EGF entry into LAMP1+ LE/Ly at 1-3 hours in wild-type (Fig 3D**, top panel**), but not *C18orf8*-deficient cells, where co-localisation remained minimal throughout the chase (**bottom panel**). As seen with EGF, *C18orf8*-deficient cells also accumulated Dil-LDL in EEA1+ EE for the duration of a 3 hours chase (Fig 3E) and trafficking of the fluid phase dye Sulforhodamine 101 (SR101) (42) into lysotracker+ LE/Ly was delayed (Fig S3F). C18orf8 therefore plays an essential role in early-to-late and/or late endosomal trafficking that precedes substrate degradation in lysosomes.

### C18orf8 is an integral component of the Mon1-Ccz1 complex

To further characterise C18orf8 function, we identified C18orf8-interacting proteins by mass spectrometry. Pull-down of N- or C-terminal HA-tagged C18orf8 revealed strong interactions with three proteins – Ccz1, Mon1A and Mon1B – the three components of the mammalian MC1 complex (Fig 4A). The interaction between overexpressed C18orf8 and endogenous Ccz1 and Mon1B was readily confirmed by immunoblot (Fig S4A). To visualise the interaction between endogenous proteins, we used CRISPR technology to knock-in a 3xMyc-tag into the C18orf8 locus, yielding an endogenous C18orf8-3xMyc fusion (Fig S4B, C). Pull-down of C18orf8-3xMyc precipitated Ccz1 and Mon1B (Fig 4B), and conversely immune precipitation of endogenous Mon1B (Fig 4B) or Ccz1 (Fig S4D) revealed the endogenous 3xMyc-tagged C18orf8. C18orf8 is composed of an N-terminal WD40 and C-terminal α-helical domain. To identify the MC1-interaction site in C18orf8, we expressed mScarlet-tagged single domains (Fig S4E). Immune precipitation revealed an interaction of Ccz1 and Mon1b with the C-terminal (AA 354-657), but not N-terminal (AA 1-362) domain of C18orf8 (Fig S4F).

**Figure 4:**
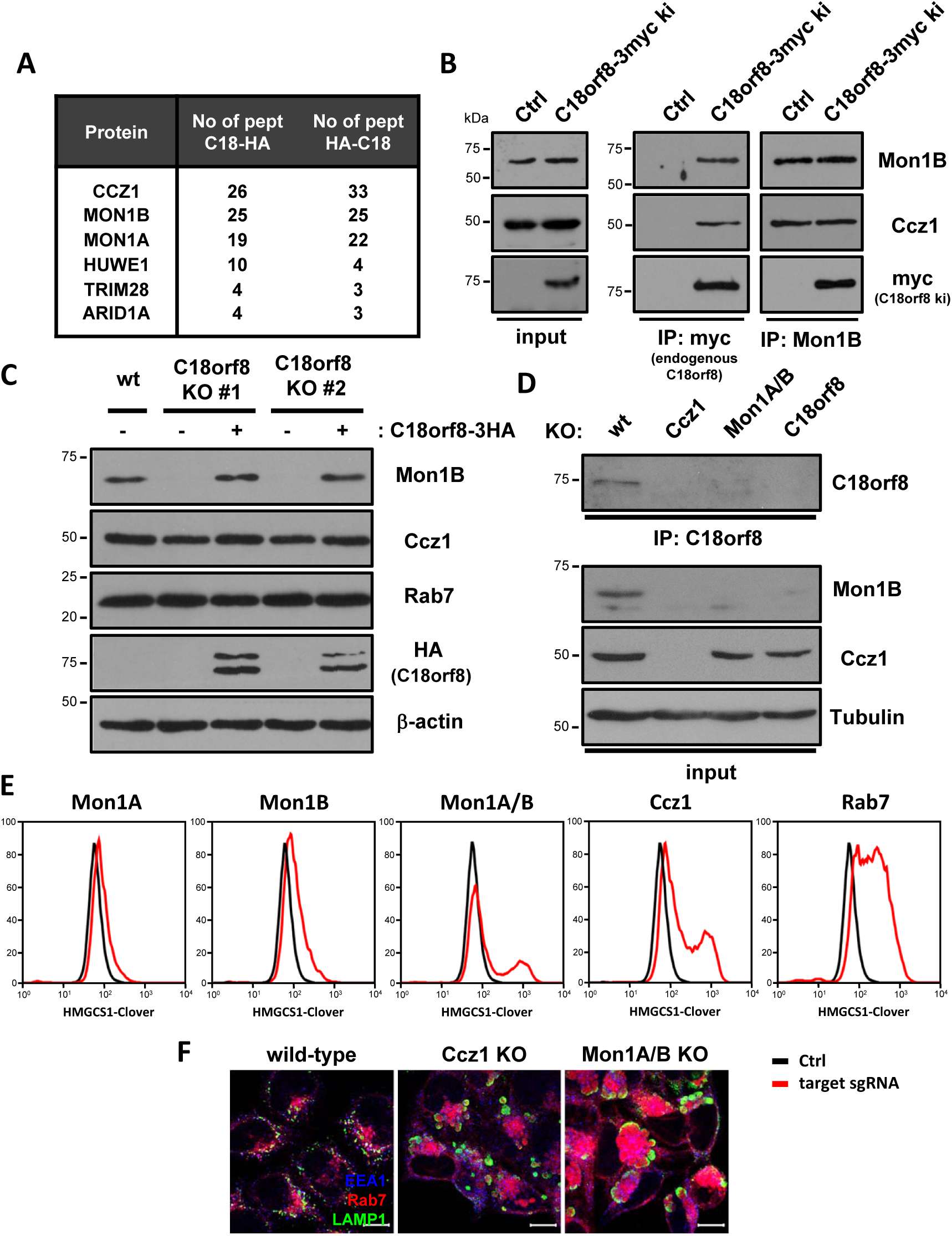
C18orf8 forms an integral component of the Mon1-Ccz1 (MC1) complex, essential for complex stability and function. (**A**, **B**) C18orf8 interacts with the Mon1-Ccz1 complex. (**A**) Immune precipitations of exogenous 3xHA-C18orf8 (N-term) or C18orf8-3xHA (C-term) were analysed by mass spectrometry. Proteins detected by >3 peptides in both C18orf8 samples and absent from control, are indicated. (**B**) Immune precipitation of an endogenous C18orf8-3xMyc fusion or Mon1B reveals a reciprocal interaction between C18orf8, Ccz1 and Mon1B (see also **Fig S4C, D**). (**C - D**) C18orf8, Mon1B and Ccz1 show reciprocal stabilisation. Immunoblot analysis of lysates from (**C**) wild-type, *C18orf8*-deficient and *C18orf8*-deficient cells complemented with C18orf8-3xHA; or (**D**) wild-type, *C18orf8-*, *Ccz1*- and *Mon1A/B*-deficient cells. For endogenous C18orf8 detection, C18orf8 was immune precipitated prior to immunoblotting (**D**). (**E - F**) *Ccz1*- and *Mon1A/B*-deficient cells show cholesterol deficiency and disruption of LE/Ly morphology. (**E**) HMGCS1-Clover CAS9 cells were transfected with sgRNAs against *Mon1A*, *Mon1B*, *Mon1A* and *Mon1B*, *Ccz1* or *Rab7*, grown for 14 days, treated overnight with mevastatin and analysed for HMGCS1-Clover expression. (**F**) Wild-type, *Ccz1*- and *Mon1A/B*-deficient cells were stained intracellularly for EEA1, Rab7 and LAMP1. Scale bars = 10µm.

The robust interaction of C18orf8 with all three MC1 components suggests C18orf8 is either a regulator or integral component of the mammalian MC1 complex. Interestingly, while *C18orf8*-deficient cells expressed normal levels of Ccz1, they showed a marked reduction in Mon1B expression (Fig 4C), which was restored by re-expression of full-length C18orf8 (Fig 4C), its C-terminal MC1-interaction domain (Fig S4F) or proteasome inhibition (Fig S4G). Mon1B expression was also decreased in *Ccz1*-deficient cells, while conversely endogenous C18orf8 expression was reduced in *Ccz1* and *Mon1A/B* double-deficient cells (Fig 4D). Mon1B stability is therefore dependent on C18orf8 and Ccz1 and in the absence of either, Mon1B is proteasomally degraded. Similarly, C18orf8 stability depends on Mon1A/B and Ccz1, whereas Ccz1 remains relatively stable in the absence of both other components (Fig 4D). We conclude that mammalian <u>M</u>on1, <u>C</u>cz1 and <u>C</u>18orf8 form a stable trimeric complex, the MCC complex, of which C18orf8 is a core component.

To determine whether the function of C18orf8 overlaps with mammalian MC1, we knocked-out *Ccz1*, *Mon1A*, *Mon1B* and *Rab7* in our *HMGCS1-Clover* reporter line. Similar to *C18orf8* depletion, loss of *Ccz1* or a combination of *Mon1A* and *Mon1B*, but neither alone, increased HMGCS1-Clover expression (Fig 4E), as did knockout of *Rab7*, a prominent hit from our screen (Fig 1E, 4E). This phenotype was restored by complementation of knockouts with their respective wild-type proteins (Fig S4I), with *Mon1A*/*B*-double-deficient cells rescued by Mon1A or Mon1B alone. Like *C18orf8*-deficient cells, cells deficient for *Ccz1*- or *Mon1A/B*-deficient contained abnormal, enlarged Rab7+ and LAMP1+ LE/Ly compartments (Fig 4F). This phenotypic similarity implies C18orf8 is required for both MCC stability and functionality. Importantly, although the C18orf8 α-helical domain is necessary and sufficient for MCC binding and Mon1B stabilization (Fig S4F), it is insufficient for full complementation of the C18orf8-deficiency phenotype which required full-length C18orf8 (**Fig S4H**). C18orf8 is therefore not simply a stabilizing factor but an active integral component of the MCC complex.

### The Mon1-Ccz1-C18orf8 complex is responsible for mammalian Rab7 activation

Yeast MC1 acts as an activating GEF for the yeast Rab7 homologue Ypt7 (28). To assess whether the mammalian MCC complex has a similar function, we expressed HA-tagged wild-type, dominant-negative (T22N) and constitutively active (Q67L) Rab7 in C18orf8-3xMyc knock-in cells and probed its interaction with endogenous MCC components. Consistent with a role in Rab7 activation, C18orf8-3xMyc, Ccz1 and Mon1B preferentially bound the inactive Rab7-T22N, but not a constitutively active Rab7-Q67L mutant (Fig 5A). Rab7 activation promotes binding and LE recruitment of its effectors RILP and ORP1L (17–20) and effector binding is commonly used to determine the activation status of Rab GTPases. In wild-type cells, an interaction between FLAG-tagged RILP and endogenous Rab7 was readily detected - indicating the presence of an active GTP-bound Rab7, whereas this interaction was lost in *C18orf8-*, *Ccz1*- and *Mon1A/B*-deficient cell lines (Fig 5B). *MCC*-deficient cells are thus unable to activate Rab7, suggesting MCC acts as a mammalian Rab7 GEF. Indeed, HA-tagged RILP was efficiently recruited to LAMP1+ LE/Ly in wild-type, but not in *C18orf8*-deficient cells, where RILP redistributed to the cytoplasm (Fig 5C). LE recruitment of the Rab7 effector ORP1L was also strongly reduced (Fig S5A). Consistent with RILP’s function in endosome mobilisation, LE motility was sharply decreased in *C18orf8*-deficient cells (Fig S5B). C18orf8, Ccz1 and Mon1A/B are therefore required for Rab7 activation and functional recruitment of the Rab7 effectors RILP and ORP1L in mammalian cells.

**Figure 5:**
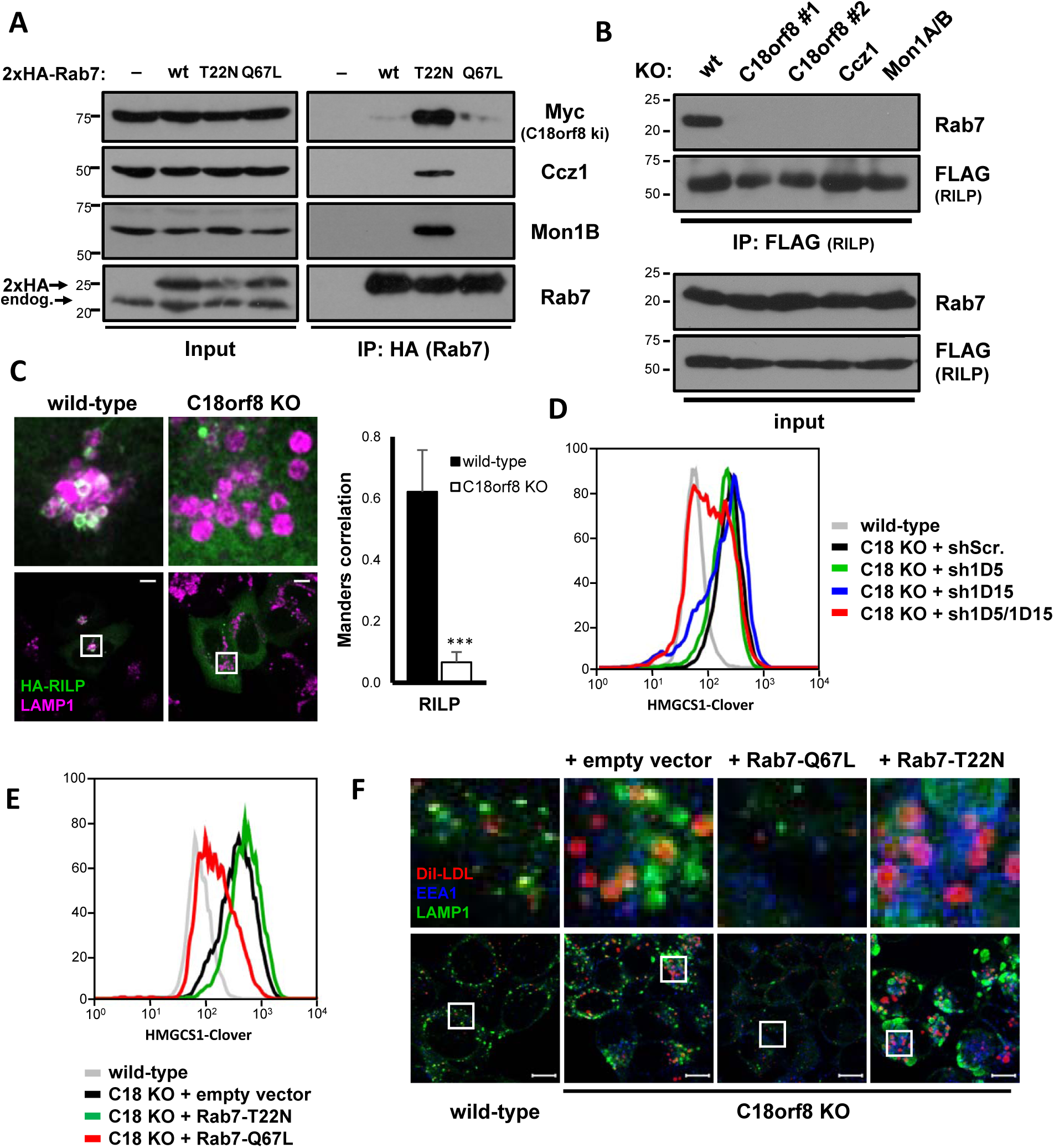
The trimeric Ccz1-Mon1-C18orf8 (MCC) complex activates mammalian Rab7. (**A**) The MCC complex binds an inactive Rab7 (T22N). Immune precipitations of 2xHA-tagged wild-type, T22N or Q67L Rab7 from *C18orf8-3xMyc* knock-in cells, were analysed by immunoblot using Myc, Ccz1 and Mon1B specific antibodies. (**B**, **C**) *C18orf8*-, *Ccz1*- and *Mon1A/B*-deficient cells lack activation-dependent recruitment of Rab7 effectors. (**B**) Immune precipitations of 3xFLAG-RILP from wild-type, *C18orf8*-, *Ccz1*- or *Mon1A/B*-deficient cells were analysed by immunoblotting for endogenous Rab7. (**C**) Wild-type and *C18orf8*-deficient cells were transfected with HA-RILP or ORP1L (**Fig S5A**) and stained intracellularly for HA (green) and LAMP1 (magenta). Manders correlation was determined for 6-10 cells from two independent experiments (*** P<0.001). (**D – F**) Cholesterol and trafficking defects in C18orf8 deficient cells can be rescued by knockdown of Rab7GAPs or expression of a constitutively active Rab7 (Q67L). (**D**) HMGCS1-Clover expression was determined at day 8 after transduction with shRNAs against TBC1D5, TBC1D15 or both. (**E**, **F**) *C18orf8*-deficient cells were transduced with 2xHA-tagged Rab7-T22N, -Q67L or an empty vector and either (**E**) analysed at day 10 by flow cytometry for HMGCS1-Clover expression, or (**F**) pulse-labelled with AlexaFluor 555 labelled EGF, incubated for 3 hours and stained intracellularly for LAMP1 and EEA1. Scale bars = 10µm.

### C18orf8 function can be by-passed by a constitutively active Rab7 or depletion of Rab7GAPs

The activation of Rab GTPases by RabGEFs is counteracted by RabGAPs that enhance intrinsic GTPase activity and revert Rab GTPases back to an inactive state. Our data suggest that LE defects in *C18orf8*-deficient cells are secondary to defective Rab7 activation. We therefore asked whether LE function could be rescued by concomitantly blocking one or more Rab7GAPs; or by the use of a constitutively active Rab7-Q67L mutant that lacks intrinsic GTPase activity. Knockdown of the Rab7GAPs TBC1D5 and TBC1D15, but not either alone, rescued HMGCS1-Clover levels in *C18orf8*-deficient cells back to wild-type (Fig 5D) and a similar rescue was obtained by expression of the constitutively active Rab7-Q67L, but not the inactive Rab7-T22N (Fig 5E). Rab7-Q67L also restored the swollen LE/Ly phenotype (Fig 5F, **green**) and Dil-LDL degradation (**red**) in *C18orf8*-deficient cells. In contrast, the T22N mutant augmented both phenotypes. The trimeric Mon1-Ccz1-C18orf8 (MCC) complex therefore acts as an activating GEF for mammalian Rab7 and its essential role in late endosomal trafficking and LDL-cholesterol uptake can be by-passed with a constitutively active Rab7 (Q67L) or by Rab7GAP depletion.

### MCC-deficient cells accumulate free cholesterol in their lysosomal compartment

How does a defect in Rab7 activation result in cellular cholesterol deficiency? *C18orf8*-deficient cells showed a severe defect in LDL trafficking (Fig 2E, 3E). We assumed this defect would cause impaired cholesterol ester (CE) hydrolysis, decreased cholesterol release and, ultimately, cellular cholesterol deficiency. To ascertain the fate of LDL-derived cholesterol, we stained our cells with Filipin III, a bacterial compound that specifically binds free cholesterol and differentiates it from CEs in LDL (43). Remarkably, *C18orf8-*deficient cells did not show a decrease in Filipin staining, but rather accumulated free cholesterol in their swollen Rab7+ and LAMP1+ LE/Ly compartment (Fig 6A, Fig 6C). Cholesterol accumulation resolved upon complementation with C18orf8-3xHA (Fig 6A) and was also observed in *Ccz1*- and *Mon1A/B*-deficient cells (Fig 6B). At the ultra-structural level cholesterol accumulation was confirmed by Theonellamides (TNM) staining (44, 45) which enriched within the enlarged MVBs of *C18orf8*-deficient cells (Fig 6D). The accumulation of free cholesterol in LE/Ly of *MCC*-deficient cells suggests that defective lysosomal cholesterol export, as opposed to LDL trafficking, is primarily responsible for the cellular cholesterol deficiency in *MCC*-deficient cells. Indeed the *MCC*-deficiency phenotype is strikingly similar to that observed in cells defective for the NPC1 cholesterol transporter (Fig 6E).

**Figure 6:**
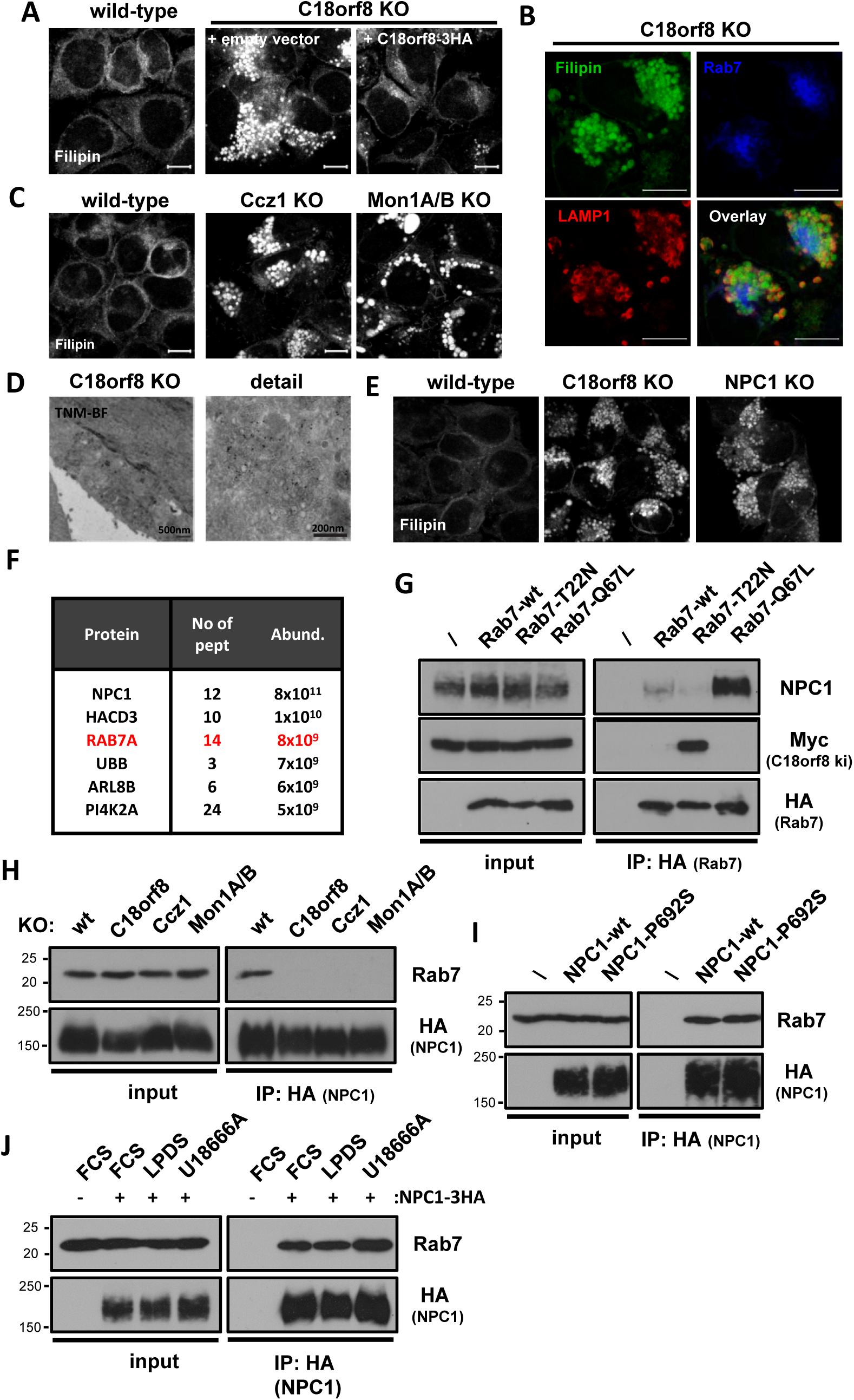
*MCC*-deficient cells lack an activation-dependent interaction between Rab7 and NPC1 and accumulate lysosomal cholesterol. (**A – C**) *MCC*-deficient cells accumulate lysosomal cholesterol and phenocopy *NPC1*-deficiency. (**A**) Filipin staining of wild-type, *C18orf8*-deficient and complemented *C18orf8*-deficient cells; or (**C**) wild-type, *Ccz1-* and *Mon1A/B*-deficient cells. (**B**) Filipin co-staining with the LE/Ly markers Rab7 and LAMP1 in *C18orf8*-deficient cells. (**D**) Theonellamides (TNM) immuno-gold labelling of *C18orf8*-deficient cells, visualised by EM. (**E**) Filipin staining of wild-type, *C18orf8*- and *NPC1*-deficient cells. Scale bars = 10µm. (**F**, **G**) Rab7 interacts with the NPC1 cholesterol transporter in activation-dependent manner. (**F**) Immune-precipitation of HA-tagged NPC1 and detection of NPC1-interacting proteins using mass spectrometry. Interaction partners detected with >2 peptides are indicated by abundance. (**G**) Immune precipitations of HA-tagged wild-type, dominant-negative (T22N) or constitutively active Rab7 (Q67L) reveal an activation-dependent interaction between Rab7 and endogenous NPC1. (**H**) The Rab7-NPC1 interaction is lost in MCC-deficient cells that lack Rab7 activation. Wild-type, C18orf8-, Ccz1 and Mon1A/B-deficient cells were stably transduced with the inactive NPC1-P692S-HA. HA-tagged NPC1 was immune precipitated and immune blotted for endogenous Rab7. The inactive NPC1-P692S was used to prevent altering lysosomal cholesterol content. (**I**, **J**) The Rab7-NPC1 interaction is independent of NPC1 activity or lysosomal cholesterol levels. (**I**) NPC1-deficient cells were complemented with HA-tagged wild-type or inactive P692S-mutant NPC1 and NPC1-HA immune-precipitations were analysed by immune blotting for endogenous Rab7. (**J**) Wild-type NPC1-HA complemented cells were treated with LPDS to decrease, or U18666A to increase lysosomal cholesterol levels and the NPC1-Rab7 interaction was probed using immune precipitation.

### Active Rab7 interacts with the NPC1 cholesterol transporter

The NPC1 transporter is critical for lysosomal export of LDL-derived cholesterol and cellular cholesterol homeostasis (Fig 1E, 1F) and *NPC1* mutations are the primary cause of Niemann Pick type C lysosomal storage disease. *C18orf8*-deficient cells show normal to elevated levels of NPC1 expression (Fig S6A), with increased NPC1 expression in *Ccz1*- and *Mon1A/B*-deficient cells (Fig S6B). In wild-type cells, NPC1 resides predominantly in the LAMP1+ Ly compartment (Fig S6C), where it co-localises with NPC2 (Fig S6D) and this localisation was unaltered in *C18orf8*-, *Ccz1*- and *Mon1A/B-*deficient cells (Fig S6C, D). Since NPC1 expression and localisation appeared normal, we used mass spectrometry to identify NPC1 interactions partners. Remarkably, among the most abundant interaction partners identified in NPC1 immune precipitations is Rab7 itself (Fig 6F). This interaction was not only confirmed in Rab7:NPC1 immune precipitations but NPC1 preferentially bound to the constitutively active Rab7-Q67L but not the inactive Rab7-T22N mutant (Fig 6G), as could be expected for a functionally important Rab7 interaction. Furthermore, consistent with an activation-dependent event, a robust interaction was observed between NPC1 and Rab7 in wild-type cells and this interaction was lost in *C18orf8-*, *Ccz1-* or *Mon1A/B-*cells that are defective in Rab7 activation (Fig 6H). The Rab7-NPC1 interaction is maintained by an inactive NPC1-P692S mutant and is therefore independent of NPC1 cholesterol export function (Fig 6I). Indeed Rab7 binding to NPC1 remains largely unaffected by treatment of cells with LPDS treatment, which decreases, and the NPC1 inhibitor U18666A which increases lysosomal cholesterol levels (Fig 6J).

### Rab7 activation by the MCC GEF drives NPC1-dependent cholesterol export

Loss of the Rab7-NPC1 interaction in *MCC*-deficient cells correlates with lysosomal cholesterol accumulation, suggesting Rab7 activation is essential for NPC1-dependent lysosomal cholesterol export. To test this hypothesis directly and avoid potential caveats created by LDL trafficking defects, we set up a lysosomal cholesterol export assay analogous to a pulse-chase. Cholesterol export was initially blocked using the reversible NPC1 inhibitor U18666A (46) to induce free cholesterol accumulation in LE/Ly (pulse). The inhibitor was then washed out in LDL-free medium (LPDS) which allowed lysosomal cholesterol efflux in the absence of further LDL-cholesterol uptake (chase). Cholesterol release from LE/Ly was monitored by Filipin staining (Fig 7A). During a 24h chase period, cholesterol was completely exported from CD63+ LE/Ly in wild-type cells, whereas export was abolished in cells deficient for the NPC1 transporter (Fig 7A, B). Similar to NPC1 deficiency, Ly cholesterol export was blocked in *C18orf8-*, *Ccz1-* and *Mon1A/B -*deficient cells in which Filipin staining remained co-localised with CD63 during the 24h chase (Fig 7A, B). To confirm that the cholesterol export defect in MCC-deficient cells depends on Rab7 activation, *C18orf8*-deficient cells were complemented with different Rab7 mutants. While the cholesterol accumulation resolved upon expression of the constitutively active Rab7-Q67L, no resolution was seen with either the wild-type or inactive Rab7-T22N (Fig 7C), confirming Rab7 dependency of the cholesterol export defect.

**Figure 7:**
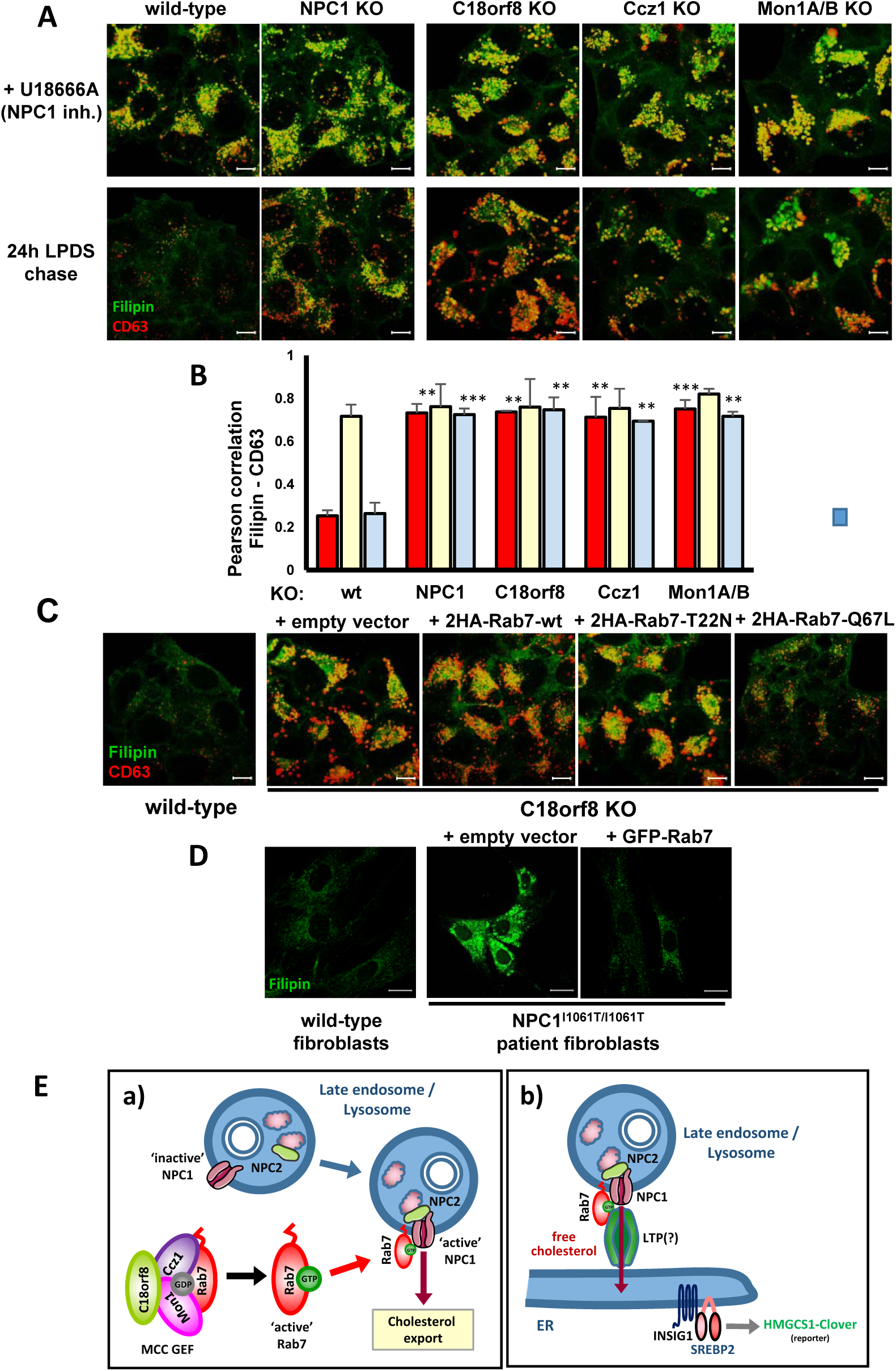
Rab7 activation by the MCC GEF controls NPC1-dependent lysosomal cholesterol export. (**A, B**) Lysosomal cholesterol export is abolished in *NPC1*-, *C18orf8*-, *Ccz1-* and *Mon1A/B*-deficient cells. (**A**) Wild-type, NPC1-, C18orf8-, Ccz1 and Mon1A/B-deficient cells were treated for 24 hours with the NPC1 inhibitor U18666A to increase lysosomal cholesterol (pulse, top panels), followed by a 24 hours chase in the presence of LPDS and mevastatin (lower panels). Lysosomal cholesterol accumulation was visualised using Filipin co-staining with the LE/Ly marker CD63. (**B**) Colocalisation of Filipin with CD63 was plotted as Pearson correlation, calculated from 3 independent experiments with 6 representative fields per experiment and >8 cells per field. ** p<0.01, *** p<0.001. (**C**) Cholesterol accumulation in C18orf8-deficient cells is abolished by overexpression of a hyperactive Rab7. C18orf8-deficient cells were transduced with a wild-type, dominant-negative (T22N) or hyperactive Rab7 (Q67L) or empty vector and co-stained with Filipin and anti-CD63. (**D**) Rab7 overexpression rescues lysosomal cholesterol accumulation in NPC patient fibroblasts. NPC1^I1061T/I1061T^ primary patient fibroblasts were transduced with a GFP-tagged wild-type Rab7 and analysed at day 7 for cholesterol accumulation using Filipin staining. Fibroblasts from a healthy individual were used as a control. Representative images are shown from 2 independent experiments. (**E**) Model for MCC and Rab7 function in lysosomal cholesterol export. The trimeric Mon1-Ccz1-C18orf8 (MCC) GEF activates mammalian Rab7, which then binds to the NPC1 cholesterol transporter and either **a**) directly activates NPC1 cholesterol export function; or **b**) assembles a down-stream membrane contact site (MCS) at which a yet-uncharacterised lipid transfer protein (LTP) mediates cholesterol transfer to the ER and/or plasma membrane. A combined Rab7 function in NPC1 activation and MCS formation would assure lysosomal cholesterol is only exported once a down-stream lipid transfer module is assembled.

NPC1 function is compromised in type C Niemann Pick (NPC) disease in which lysosomal cholesterol accumulates in multiple organs. The common NPC1^I1061T^ mutation destabilizes the NPC1 transporter, resulting in its ER retention, and subsequent degradation (47). However, any transporter reaching its correct lysosomal location remains functional (48). We asked whether increased Rab7 expression improves NPC1 function, promoting clearance of lysosomal cholesterol in mutant NPC1 fibroblasts. Remarkably, lentiviral overexpression of a GFP-tagged wild-type Rab7 strongly reduced cholesterol accumulation in NPC1^I1061T/I1061T^ patient fibroblasts (Fig 7D), suggesting Rab7 activity is limiting for lysosomal cholesterol export in NPC disease.

Taken together our results show that Rab7 and its trimeric Mon1-Ccz1-C18orf8 (MCC) GEF control cellular cholesterol homeostasis through coordinated regulation of LDL trafficking and NPC1-dependent lysosomal cholesterol export, making it a potential therapeutic target in Niemann Pick disease.

## Discussion

The endocytic uptake of LDL-cholesterol is a major source of cellular cholesterol. Using an endogenous cholesterol sensor, HMGCS1-Clover, we screened for and identified >70 genes involved in the cellular LDL-cholesterol uptake pathway. Of these, we characterised C18orf8 as a novel core component of the Mon1-Ccz1 (MC1) GEF for mammalian Rab7. *C18orf8*-deficient cells are defective Rab7 activation, resulting in impaired LDL trafficking and cellular cholesterol depletion. The unexpected accumulation of free cholesterol in the swollen LE/Ly compartment of *C18orf8*-deficient cells, suggested an additional critical role for Rab7 in cholesterol export out of lysosomes. We show that active Rab7 binds the NPC1 cholesterol transporter and licenses lysosomal cholesterol export. This pathway is defective in *C18orf8*-, *Ccz1*- and *Mon1A/B*-deficient cells and restored by a constitutively active Rab7 (Q67L). The trimeric <u>M</u>on1-<u>C</u>cz1-<u>C</u>18orf8 (MCC) complex thus plays a central role in cellular LDL-cholesterol uptake, coordinating Rab7 activation, LDL trafficking and NPC1-dependent lysosomal cholesterol export.

### A CRISPR screen provides a comprehensive overview of trafficking factors required for cellular LDL-cholesterol import

Genome-wide CRISPR screens provide a powerful tool to interrogate intracellular pathways. To avoid artefacts associated with exogenous reporters, we generated an endogenous SREBP2 knock-in reporter, HMGCS1-Clover, and screened for genes regulating cellular cholesterol homeostasis. The predominant pathway identified by our screen is LDL-cholesterol import, as HeLa cells, like many tissue-culture cells, rely predominantly on cholesterol import rather than *de novo* biosynthesis. The screen identified multiple stages in the LDLR-LDL trafficking pathway (Fig 1E, F), including **i)** LDLR folding and trafficking to the cell surface; **ii)** endocytosis of the LDLR-LDL complex; **iii)** LDL trafficking to the LE/Ly compartment; and **iv)** NPC1 dependent cholesterol export from LE/Ly. Functional complexes identified include COG, Exocyst, AP2, V-ATPase, PI3K, PIKfyve, CORVET, HOPS, FERARI and GARP; poorly characterised genes include the chaperone Clusterin (also known as Apolipoprotein J) and the metalloprotease ADAM28. Our functional genetic approach therefore reveals a comprehensive genome-wide analysis of essential cellular components required for LDL-cholesterol import, and identifies novel candidate genes in this pathway.

### C18orf8 is a core component of the Mon1-Ccz1-C18orf8 (MCC) GEF for mammalian Rab7

Spatial and temporal control of Rab7 activation is critical to coordinate substrate trafficking and degradation in LE/Ly (16). In yeast the dimeric Mon1-Ccz1 (MC1) GEF activates the Rab7 homologue Ypt7 (27, 28) and a similar function has been suggested for mammalian MC1 (35, 36). We find that a stable trimeric complex of Mon1A/B, Ccz1 and C18orf8 (MCC) is required for Rab7 activation in mammalian cells, as defined by effector recruitment. *C18orf8*, *Mon1* and *Ccz1* are conserved from animals to plants, but a *C18orf8* orthologue is missing from laboratory yeast and most other fungi, either due to divergence of the endocytic machinery or because C18orf8 function is incorporated into other components.

Rab7 is a key regulator of LE homeostasis and *C18orf8-*, *Ccz1-* and *Mon1A/B*-deficient cells show a broad range of LE defects including LE morphology, endosomal substrate trafficking, lysosomal degradation and cholesterol export. Consistent with a previous Rab7 depletion study (49), *C18orf8*-deficient cells accumulate enlarged MVBs; and consistent with current Rab5-to-Rab7 conversion models (32) we observe a delay in early-to-late endosomal trafficking. Whereas LDL trafficking was previously suggested to involve only Mon1B (50), we find that only the combined depletion of Mon1A/B causes cholesterol deficiency, suggesting redundancy of Mon1 homologues.

Besides LE trafficking, Rab7 is an important factor in autophagy (51). During preparation of this manuscript, a role for C18orf8 was reported as a regulator of autophagic flux (52). Like us, this group reports a robust interaction between C18orf8 and mammalian MC1. Without direct evidence of Rab7 activation, C18orf8 is however postulated to be a MC1 regulator. We find C18orf8 is not a regulator, but a core component of the trimeric mammalian MCC complex, which acts as a Rab7 GEF. Stability of the three core components is interlinked, with Mon1B stability dependent on C18orf8 and Ccz1, and C18orf8 stability dependent on Ccz1 and Mon1B. Ccz1 appears relatively stable in the absence of the other components and may therefore form a scaffold for Mon1A/B and C18orf8 assembly. The intricate relationship between C18orf8 and Mon1A/B is intriguing as Phyre2 prediction suggests C18orf8 adopts a clathrin-like fold, while Mon1A/B shows structural similarity to the AP2μ subunit (53). Whether this structural similarity has functional consequences, remains to be established.

### What is the role of C18orf8 within the MCC GEF complex?

The structural interdependence of the MCC subunits complicates *in vivo* studies to assess the direct role of C18orf8 in mammalian MCC. Yeast Ccz1p and Mon1p are necessary and sufficient for Ypt7 binding and GDP-to-GTP exchange, with most of the MC1-Ypt7 binding interface mediated by Mon1p (28, 34). Residues critical for Ypt7 binding are conserved in mammalian MC1 and a dimeric complex of mammalian Mon1A and Ccz1 was shown to have limited *in vitro* Rab7 GEF activity (36). While this complex might have retained residual C18orf8 due to its isolation from mammalian cells, future studies will address whether C18orf8 contributes towards *in vitro* MC1 GEF activity.

*In vivo,* C18orf8 function goes beyond being a scaffold for MCC stabilisation. The C18orf8 C-terminal α-helical domain is sufficient to bind MC1 and stabilize Mon1B, but insufficient to fully restore MCC function (Fig S4F, I). The C18orf8 N-terminus thus has an independent function and may bind other trafficking components and/or membrane lipids. The timing of Rab7 activation is strictly regulated to coordinate Rab5-to-Rab7 conversion and likely requires control of MCC by upstream components (32). Membrane association allosterically activates yeast MC1 and increases its GEF activity ∼1600-fold (29). Like yeast Mon1p and Ccz1p, mammalian Mon1A binds PtdIns3P and PtdSer (27), while positively charged residues in the N-terminal β-propeller of C18orf8 are well-positioned to bind negatively charged phospholipids. In analogy to AP2 (54), PtdIns binding might open the MCC complex towards Rab7 binding, thus allosterically activating MCC upon membrane association.

### Rab7 controls NPC1-dependent cholesterol export from late endosomes

Cholesterol is hydrolysed from CEs in a LE/Ly compartment, and NPC1 is essential for its subsequent export to the ER, mitochondria and plasma membrane. Increased expression of our SREBP2 reporter indicates *C18orf8*-, *Ccz1*- and *Mon1A/B*-deficient cells are starved of ER cholesterol, yet paradoxically they accumulate free cholesterol in a swollen LE/Ly compartment. This suggested a novel role for Rab7 in lysosomal cholesterol export. We find an active Rab7 binds the lysosomal cholesterol transporter NPC1 and licenses its export of LDL-derived cholesterol from LE/Ly. The inability of MCC-deficient cells to activate Rab7, results in a Niemann-Pick-like phenotype, even though NPC1/NPC2 expression and localisation are intact. Rab7 therefore functions as a novel regulator of NPC1 and its export of lysosomal cholesterol.

The mechanism by which Rab7 controls NPC1-dependent lysosomal cholesterol export remains unclear. Rab7 may **(i)** directly activate the NPC1 transporter, thus licensing lysosomal cholesterol export (Fig 7E**, panel a**); **(ii)** assemble a lipid transfer hub down-stream of NPC1 to allow inter-organelle cholesterol transfer at membrane contact sites (MCS) (**panel b**); or **(iii)** act by a combination of both.

### Activation of the NPC1 transporter by Rab7

The activity of many transporters is tightly regulated, for instance the lysosomal calcium channel TRPML1 is specifically activated by PtdIns(3, 5)P_2_ at LE/Ly (55). Even so, the NPC1 transporter is believed to be constitutively active and to date no NPC1 regulators have been identified. So why would Rab7 regulate NPC1 activity? Inappropriate cholesterol export by a nascent NPC1 could harm the composition and functionality of non-lysosomal compartments. To prevent cholesterol export outside LE/Ly, the interaction of cholesterol-loaded NPC2 with NPC1 depends on the acidic lysosomal pH (7). Activation of NPC1-dependent cholesterol export by Rab7 might further restrict cholesterol transfer to LE/Ly. The Rab7-NPC1 interaction is independent of lysosomal cholesterol content. Even so, variations in Rab7 activity might tune lysosomal cholesterol export to cellular or organelle-specific conditions, such as MCS formation or organelle division. Future studies will aim to identify the cytoplasmic Rab7 binding site on NPC1 and its putative function in regulating NPC1 activity.

### A role for Rab7 in cholesterol transfer at membrane contact sites

Besides NPC1 activation, Rab7 might control lysosomal cholesterol export through the formation of inter-organelle membrane contact sites (MCS) and/or the recruitment of lipid transfer proteins (LTPs) down-stream of NPC1. At least three Rab7 effectors - ORP1L, Protrudin (ZFYVE27) and PDZD8 - are involved in ER-to-LE MCS formation (19, 20, 56, 57) and Rab7 itself has been implicated in ER-to-mitochondria MCS formation (58). MCS are of key importance for rapid inter-organelle lipid exchange, yet whether Rab7-mediated MCS are involved in cholesterol transfer remains controversial. The Rab7 effector ORP1L was reported to act as a LTP linking the ER and LE/Ly and transporting cholesterol towards the ER in a PtdIns(4, 5)P_2_ / PtdIns(3, 4)P_2_-dependent manner (12, 59). Its depletion however only causes a minor steady-state change in lysosomal cholesterol, as does depletion of other LTPs and tethering proteins implicated, including STARD3, ORP5 and SYT7 (13, 14, 60). This suggests redundancy between LTP(s) or an uncharacterised LTP required for lysosomal cholesterol export. In contrast, our *C18orf8*-, *Ccz1-* and *Mon1A/B*-deficient cells are entirely defective in lysosomal cholesterol export and fully phenocopy an NPC1-deficiency phenotype. The LTP(s) responsible for lysosomal cholesterol export may therefore be Rab7 effectors. Future studies will aim to identify the Rab7-dependent LTP(s) responsible for lysosomal cholesterol accumulation in *MCC*-deficient cells.

### The MCC Rab7 GEF coordinates LDL trafficking and lysosomal cholesterol export

LDL-cholesterol uptake is a highly efficient process, with each LDLR molecule estimated to endocytose one LDL particle containing ∼1600 CE/cholesterol molecules every 10min (3). Down-stream LDL processing must be tightly coordinated to prevent a backlog in the endocytic pathway. CE hydrolysis therefore likely commences *en route* to NPC1+ Ly to allow the large LDL particle to unpack; whereas the arrival and export of cholesterol in Ly must be linked to the formation and transport capacity of MCS. As a central regulator or LE homeostasis, Rab7 is well suited to coordinate these processes. Indeed, *MCC*-deficient cells show defects in LDL trafficking and lysosomal cholesterol export, indicating both are linked to the Rab7 nucleotide cycle. We propose that Rab7 and its MCC GEF are central regulators of cellular LDL-cholesterol uptake, coordinating LDL trafficking, NPC1-dependent lysosomal cholesterol export and lipid transfer at MCS. Besides cholesterol, the Rab7 lipid transfer hub might mediate lysosomal egress of other lipids, with VPS13A/C prominent Rab7 effectors (61, 62).

### Rab7 functionality in Niemann Pick disease

Mutations in NPC1 and NPC2 cause the lethal Niemann Pick type C (NPC) lysosomal storage disease. Disease-associated mutations commonly destabilize NPC1, leading to its ER retention and proteasome- or lysosome-mediated degradation (47, 63). The most prevalent NPC1 mutation, I1061T, destabilizes NPC1, yet the mutant protein retains its cholesterol export function and the disease phenotype can be restored by overexpression of the mutant NPC1 (48). Confirming earlier observations (64), we show lentiviral overexpression of a GFP-tagged wild-type Rab7 also reduces cholesterol accumulation in NPC1^I10161T/I1061T^ patient-derived primary fibroblasts (Fig 7D). Whereas this rescue was previously ascribed to enhanced vesicular trafficking, we suggest that Rab7 overexpression directly enhances NPC1-dependent lysosomal cholesterol export. This effect depends on an active Rab7 and is not observed with the inactive Rab7-T22N, whereas the hyperactive Rab7-Q67L shows an intermediate phenotype (Fig S7). A modest increase in Rab7 activity therefore increases lysosomal cholesterol clearance in NPC1-mutant cells while optimal clearance requires both Rab7 activation and inactivation in a time- and location-dependent manner.

In summary, we have identified a novel function for the Rab7 GTPase and its trimeric Mon1-Ccz1-C18orf8 (MCC) GEF in the coordination of late endosomal LDL trafficking and NPC1-dependent lysosomal cholesterol export. Future studies will aim to further characterise the mechanism behind Rab7 function in lysosomal cholesterol export and explore its therapeutic potential in Niemann Pick disease.

## Acknowledgments

We are grateful to the following people for their help in this study: Dr. Michael Bassik (Stanford University) for kind donation of the genome-wide CRISPR/Cas9 sgRNA library and Prof. Ron Kopito and Dr. Matthew Porteus (Stanford University) for the pDonor CRISPR knock-in plasmid. FACS experiments were enabled by Dr. Reinard Schulte and the CIMR FACS core facility team and Confocal microscopy by Matthew Gratian and Mark Bowen. Stuart Bloor kindly assisted with the Illumina sequencing, Dr. James Williamson analysed samples by mass spectrometry and Dr. Nick Matheson critically reviewed the manuscript. Special thanks to Zhengzheng Sophia Liang for fruitful discussions and moral support. This work was financially supported by a Wellcome Trust Principal Research Fellowship to P.J.L (084957/Z/08/Z). JPL was supported by MRC research grant MR/R0009015/1. The Cambridge Institute for Medical Research (CIMR) was in receipt of a Wellcome Trust strategic award (100140). The authors declare no competing financial interests.

## Author contributions

DJHB and PJL conceived the project. Experiments were carried out by DJHB, AS, IB, MLMJ and DE. DJHB analysed the data and prepared the figures. DJHB and PJL wrote the manuscript. JJCN and JPL advised on the project and critically reviewed the manuscript.

## Materials and Methods

### Plasmids

pHRSIN.pSFFV and pHRSIN.pCMV lentiviral expression constructs harbouring pGK Puromycin, pGK Hygromycin or pSV40 Blasticidin resistance have been reported previously (65). The C18orf8 open reading frame was cloned from HeLa cell cDNA, modified with an N- or C-terminal 3xHA-tag and inserted into pHRSIN.pSFFV pGK Hygro and pHRSIN.pSFFV pGK Puro respectively. Ccz1, Mon1A and Mon1B were cloned from IMAGE clones (Dharmacon), modified with a C-terminal 3xMyc tag and inserted into pHRSIN.pSFFV pGK Hygro. Mon1A was corrected by PCR to add a missing N-terminus. 2xHA-Rab7 constructs were cloned from GFP-HA-Rab7 wild-type, T22N and Q67L constructs (Addgene #28047, #28048, #28049, created by Prof. Qing Zhong, University of California Berkeley, USA (21)) and inserted into pHRSIN.pCMV pGK Puro. HA-RILP and HA-ORP1L constructs have been described previously (66, 67). RILP was recloned with an N-terminal 3xFLAG tag and inserted into pHRSIN.pSFFV pGK Puro. FAM160B2 was amplified from human stem cell cDNA and inserted in pHRSIN.pSFFV pGK Puro with a C-terminal 3xHA-tag.

The pDonor vector used for C-terminal knock-in was a kind gift from Prof. Ron Kopito and Dr. Matthew Porteus (Stanford University, USA) and modified with Clover (Addgene #49089, created by Prof. Kurt Beam, University of Colarado (38)) or a 3xMyc C-terminal tag replacing the original TAP-tag. 750-800bp flanking arms for HMGCS1 or C18orf8 were cloned from HeLa cell genomic DNA and inserted into pDonor using Gibson Assembly (NEB), yielding pDonor HMGCS1-Clover Puro and pDonor C18orf8-3xMyc Puro respectively. To allow selection for multiple-allele insertion, the puromycin resistance cassette of pDonor was replaced with hygromycin or blasticidin resistance cassettes, yielding pDonor HMGCS1-Clover Hygro and pDonor HMGCS1-Clover Blast respectively.

The pHRSIN.pSFFV 3xFLAG-NLS-CAS9-NLS pSV40 Blast construct used for stable CAS9 expression was described previously (68). The genome-wide Bassik sgRNA library (10 sgRNAs per gene, total 220,000 sgRNAs) was a kind gift from Dr. Mike Basssik (Stanford University, USA, (39)). sgRNAs for individual genes were cloned into a modified pKLV.U6 pGK Puro-2A-BFP vector originally created by Dr. Kosuke Yusa (Wellcome Sanger Institute, Hinxton, UK).

sgRNA sequences used were: SREBF1 sg1 GGTCACAGTGGTCGTTACAG, sg2 CCTGTAGAGAAGCCTCCCGG; SREBF2 sg1 AGTGCAACGGTCATTCACCC, sg2 CTCACCGTCGATGTCTCCCA, sg3 AGCCGGGCGATGGACGACAG; LDLR sg1 CGGCGAATGCATCACCC, NPC1 sg1 CGTCAGCGTCCTTCCCACAC; AP2M1 sg1 CCTGTCTCGGAATTCTGT; ANKFY1 sg1 TATTCGCTTCTACCAGA, sg2 TAGCGGGTACTCTGTCT, sg3 ACGCAGCCAAACGCCTGTAC, sg4 GTACAGGCGTTTGGCTGCGT; VPS16 sg1 GCAGTGGAAGAGTGGACCCG; ZFYVE20 sg1 CTCAGCTCTGAATAATCGGG; INSIG1 sg1 CGTAGCTAGAAAAGCTA, sg2 CTGCTGTCCCGCAGCAGGG, sg3 CAGCCCCTACCCCAACACC, sg4 TCAACCTGCTGCAGATCCAG; C18orf8 sg1 GCGGCCGGTGCAGTTCGAGA, sg2 TTCCGAAAGAGACATCGCAA; Mon1A sg1 GTTGGGGCCCGTGTAGTAAA; Mon1B sg1 GCAGGTATAGAGCTCGAATT; Ccz1 sg1 AATGAGAAGATTAGAAATGT; Rab7A sg1 AGGCGTTCCAGACGATTGCA; β2m sg1 GGCCGAGATGTCTCGCTCCG.

shRNAs were cloned into the lentiviral pHRSiren vector harbouring puromycin or hygromycin resistance cassettes. shRNA sequences used were: TBC1D5 sh1 GAAGCCATATCGCAGAGCTA, TBC1D15 sh1 AATGGGACATGGTTAATACAGTT, non-targeting sh1 GGGTATCGACGATTACAAA.

### Antibodies

Primary antibodies used include: mouse anti-HA-tag (16B12, Biolegend, #901501), rabbit anti-HA-tag (Poly9023, Biolegend, #923501), rat anti-HA-tag (3F10, Sigma-Aldrich, **#ROAHAHA),** rabbit anti-Myc-tag (71D10, Cell Signalling, #2278), mouse anti-Myc-tag (9B11, Cell Signalling, #2276), mouse anti-FLAG-tag (M2, Sigma-Aldrich, #F1804), rabbit anti-GFP-tag (Thermo-Fisher, #A11122), rabbit anti-Rab5 (C8B1, Cell Signalling, #3547), rabbit anti-Rab7 (D95F2, Cell Signalling, #9367), rabbit anti-Rab7 (EPR7589) AlexaFluor647 conjugate (Abcam, #ab198337), mouse anti-C18orf8 (OTI4E4, Novus, #NBP2-01949), rabbit anti-C18orf8 (Proteintech, #20111-1-AP), rabbit anti-Mon1B (Novus, #NBP1-92131), rabbit anti-Mon1B (Proteintech, #17638-1-AP), mouse anti-Ccz1 (B-7, Santa Cruz, #sc-514290), mouse anti-EEA1 (14/EEA1, BD Biosciences, #610456), mouse anti-LAMP1 (H4A3, Biolegend, #328601), mouse anti-LAMP1 (H4A3) BV421 conjugate (BD Biosciences, #562623), mouse anti-LAMP1 (H4A3) AlexaFluor647 conjugate (Biolegend, #121610), mouse anti-CD63 (MEM-259, Thermo-Fisher, #MA1-19281), mouse anti-HMGCS1 (A6, Santa Cruz, #sc-166763), mouse anti-HMGCR (C1, Santa Cruz, #sc-271595), mouse anti-LDLR (C7) PE-conjugate (BD Biosciences, #565653), Mouse anti-alpha-tubulin (DM1A, Thermo-Fisher, #62204), mouse anti-beta-actin (AC-74, Sigma-Aldrich, #A5316), rabbit anti-BODIPY-FL (Thermo-Fisher, #A5770).

Secondary antibodies used include: goat-anti-rabbit IgG (H+L) AlexaFluor546 and AlexaFluorPlus647 (Thermo-Fisher, #A11035, #A32733), goat-anti-mouse IgG (H+L) AlexaFluor568 and AlexaFluor647 (Thermo-Fisher, #A11031, #A21236), goat-anti-mouse IgG1 AlexaFluor546 (Thermo-Fisher, #A21123), donkey anti-rat IgG (H+L) CF568 (Biotium, # 20092), donkey anti-mouse IgG (H+L) AlexaFluor647 (Thermo-Fisher #A31571), goat-anti-mouse IgG2a BV421 (Jackson Immunoresearch, #115-675-206), goat-anti-mouse (H+L) HRP-conjugate (Jackson Immunoresearch, #115-035-146), goat-anti-rabbit (H+L) HRP-conjugate (Jackson Immunoresearch, #111-035-144), Mouse TrueBlot ULTRA HRP-conjugate (Rockland, #18-8817-33).

For immune precipitation: EZview Red anti-HA, anti-c-Myc and anti-FLAG M2 Affinity Gel (Sigma-Aldrich, #E6779, #E6654, #F2426). Protein A and Protein G-Sepharose (Sigma-Aldrich, #P3391, #P3296), IgG sepharose 6 Fast Flow (#GE17-0969-01).

### Cell lines

HeLa and 293T cells were maintained respectively in RPMI-1640 and IMDM (Sigma-Aldrich) supplemented with 10% fetal calf serum (FCS). For sterol depletion cells were washed in PBS and incubated overnight in RPMI supplement with 5% lipoprotein-depleted serum (LPDS, Biosera), 10μM mevastatin (Sigma-Aldrich) and 50μM mevalonate (Sigma-Aldrich). NPC1^I1061t/I1061T^ patient-derived primary fibroblasts (GM18453) and healthy controls (GM08399) were obtained from Coriell Institute for Medical Research (New Jersey, USA) and grown in MEM with Earl’s salt (Sigma-Aldrich) supplemented with 15% FCS and non-essential amino acids (Gibco).

CRISPR knock-out lines were created by transient transfection of HeLa cells using TransIT HeLa Monster (Mirus). Transfected cells were selected on puromycin 24-72h post-transfection and single-cell cloned >7 days post-transfection. Knock-out clones were characterised by flow cytometry and Western blotting.

Stable protein overexpression was achieved using lentiviral transduction. Briefly, 293T cells were co-transfected in a 1:1 ratio with a lentiviral expression vector (pHRSIN/pHRSiren/pKLV) and the packaging vectors pMD.G and pCMVR8.91 using TransIT-293 (Mirus). Supernatant was harvested at 48h post-transfection and transferred onto target cells. Cells were spun 45min at 1800rpm to enhance infectivity and incubated with virus overnight. Transduced cells were selected for stable transgene expression with appropriate antibiotics from 48h post-transduction.

### CRISPR knock-in of C-terminal tags

HeLa cells were transiently transfected with sgRNA, CAS9 and pDonor vector using TransIT HeLa Monster (Mirus). To create a C18orf8-3xMyc knock-in a single donor vector (pDonor C18orf8-3xMyc Hygro) was used, whereas for HMGCS1-Clover knock-in a combination of three donor plasmids (pDonor HMGCS1-Clover Puro, Hygro, Blast) was used to select for multiple-allele integration. Transfected cells were transiently selected for sgRNA expression using puromycin at 24-72h post-transfection, followed by selection of the knock-in cassette using hygromycin or a combination of puromycin, hygromycin and blasticidin at 7 days post-transfection. Selected cells were transiently transfected with Cre recombinase to remove resistance cassettes and single-cell cloned. Single cell clones were characterised using flow cytometry and Western blotting.

### Genome-wide CRISPR screening

HeLa HMGCS1-Clover cells were stably transduced with Cas9. CRISPR sgRNA library lentivirus was produced as indicated above and titrated on HeLa HMGCS1-Clover/Cas9 cells. For screening, 1×10^8^ cells were transduced at 30% infectivity (>100-fold coverage), puromycin selected, grown for 9 days and sorted by FACS for a HMGCS1-Clover^high^ phenotype. Sorted cells were grown for another 9 days and sorted a second time for the same phenotype. Cells were finally grown for another 5 days, harvested and genomic DNA was extracted using a Genra Puregene Core kit A (Qiagen). Integrated sgRNAs were amplified via two rounds of PCR, using the following primer sets: outer 5’-AGGCTTGGATTTCTATAACTTCGTATAGCATACATTATAC-3’ and 5’-ACATGCATGGCGGTAATACGGTTATC-3’; inner 5’-AATGATACGGCGACCACCGAGATCTACACTCTCTTGTGGAAAGGACGAAACACCG-3’ and 5’-CAAGCAGAAGACGGCATACGAGATnnnnnnnGTGACTGGAGTTCAGACGTGTGCTCTTCCGATCCGACTCGGTGCCACTTTTTC-3’. The second (inner) PCR introduced adaptors for Illumina sequencing, with nnnnnnn indicating variable index sequences. PCR products were purified using AMPure XP beads (Agencourt), quantified on a DNA-1000 chip (Agilent) and sequenced on a Miniseq sequencer (Illumina). sgRNA abundance in the sorted sample was compared against a library sample of similar age (68) and sgRNA enrichment was calculated using the RSA algorithm under default settings (69). Hits with p<10^-5^ were manually annotated into function pathways (cholesterol import, endocytosis, endosomal trafficking, ER/Golgi trafficking, protein folding/glycosylation, SREBP pathway) (Fig 2C, Fig 3). The full dataset is available in **Table S1**.

### Metabolic labelling and pulse-chase

Cells were sterol-depleted overnight, starved for 30 min at 37°C in methionine-free, cysteine-free RPMI containing 5% dialysed LPDS, labelled with [^35^S]methionine/[^35^S]cysteine) (Amersham) for 10 min and then chased in RPMI with 10% FCS supplemented with 2 µg/ml 25-hydroxy-cholesterol and 20 µg/ml cholesterol. Samples taken at the indicated time-points were lysed in 1% Digitonin/TBS as above. Immunoprecipitations were performed as above, washed with 1% Tx-100/TBS and samples separated by SDS-PAGE and processed for autoradiography with a Cyclone scanner (Perkin-Elmer).

### Immunoblotting and Immunoprecipitation

For immunoblotting, cells were lysed in 1% IGEPAL (Sigma-Aldrich) in TBS pH 7.4 with Roche complete protease inhibitor or in 2% SDS in 50mM Tris pH7.4 in the presence of Benzonase (Sigma-Aldrich). Post-nuclear supernatants were heated at 70°C in SDS sample buffer, separated by SDS-PAGE and transferred to PVDF membranes (Millipore). Membranes were probed with the indicated antibodies and reactive bands visualised with ECL, Supersignal West Pico or West Dura (Thermo Scientific).

For immunoprecipations, cells were lysed in 1% digitonin, 1% IGEPAL or 0.5% IGEPAL in TBS pH 7.4 with Roche protease inhibitor. Samples were precleared with protein A/IgG-Sepharose and incubated with primary antibody and protein A/protein G-Sepharose or antibody conjugated agarose for at least 2 hrs. After 5 washes in 0.2% detergent, proteins were eluted in 2% SDS, 50mM Tris pH7.4, separated by SDS-PAGE and immunoblotted as described. For RILP/Rab7 immune precipitations, lysis buffers were supplemented with 10mM MgCl_2_ and 1mM EDTA.

### Flow cytometry

Cells were washed in PBS, detached by trypsinising for 5 min, pelleted and where necessary antibody stained for 30min on ice. After washing in ice-cold PBS, cells were analysed on a FACS Calibur or Fortessa (BD Biosciences) and analysed in FlowJo.

### Immune fluorescence

Cells were grown on 10mm coverslips, fixed for 15 min in 4% formaldehyde (Polysciences), washed 3 times with PBS and incubated 10 min with 15mM Glycine. Cells were permeabilised for 1 hour with 0.05% saponin, 5% goat serum in PBS and stained for 1-2 hours with primary antibody as indicated in 3% BSA, 0.05% saponin, PBS. Coverslips were washed 3 times 5 min in 0.1% BSA, 0.05% saponin, PBS and incubated 1h with fluorochrome-conjugated secondary, washed 3 more times and embedded in Prolong Antifade Gold (Thermo Scientific). Stainings with Rab5 and Rab7 antibody (Cell Signalling) antibody were performed overnight in 3% BSA, 0.3% Tx-100, PBS at 4°C, followed by staining with additional primary antibody and secondary antibodies as above.

For Dil-LDL co-staining, cells were permeabilised for 40 sec in 0.01% digitonin (Merck) and stained in 3% BSA without detergent. For Filipin staining, cells were fixed and incubated 1 hour with 0.05mg/ml Filipin III (Cayman) in 0.5% BSA. For Filipin co-staining, primary and secondary antibody stainings were performed in the presence of Filipin and without detergent. Fluorescent staining was recorded on a Zeiss LSM880 Confocal microscope using a 63x oil objective and analysed using Zen software (Zeiss).

For ORP1L/RILP-LAMP1 co-localisation, cells were transiently transfected with HA-tagged RILP or ORP1L constructs using Effectine (Qiagen), harvested at 24 hours post-transfection and fixed in 3.7% formaldehyde in PBS for 15-20 min. After 10min permeabilized with 0.1% TritonX-100 in PBS, coverslips were stained with HA and LAMP1 primary antibodies and donkey anti-rat CF568 and donkey anti-mouse Alexa647 secondary antibodies and mounted. Samples were imaged on a Leica SP8 microscope adapted with a HCX PL 63x 1.32 oil objective, solid-state lasers, and HyD detectors. Colocalization was reported as Mander’s coefficient calculated using JACoP plug-in for ImageJ on the basis of 2 independent experiments.

### LDL and EGF trafficking assays

For LDL trafficking assays, cells were starved 30min in RPMI with 5% LPDS at 37°C, labelled 15 min at 4°C with 20 μg/ml Dil-LDL in RPMI with 5% LPDS and 10mM HEPES, washed and chased for indicated times in full medium. For EGF trafficking, cells were starved for 1 hour in HBSS (Gibco), incubated with 100ng/ml EGF-AlexaFluor 555 for 5 min at 4°C, washed and chased in full medium. At indicated times, cells fixed in 4% formaldehyde and either mounted directly or stained for indicated markers as described above. Fluorescent staining was recorded using a Zeiss LSM880 Confocal microscope with a 63x oil objective. EEA1+ and LAMP1+ vesicles were annotated in an automated fashion using Volocity software (Perkin-Elmer) and the percentage EGF within these vesicles was determined as a percentage of total EGF fluorescence for 5-6 fields per sample with >8 cells per field. To include luminal content of enlarged LAMP1+ vesicles area filling function was enabled and enlarged vesicles were trimmed back by one pickle to prevent artificial area merging in areas of high vesicle density. Statistical significance was determined using two-tailed Student’s T-Test with **p<0.01 and ***p<0.001 and error bars reflecting standard error of mean (SEM).

### Lysosomal cholesterol export assay

Cells were seeded on coverslips and treated for 24 hours with 2 μM U18666A (Santa Cruz) in RPMI with 10% FCS, washed with PBS and either fixed directly with 4% paraformaldehyde (pulse); or incubated for 24 hours chase in RPMI with 5% LPDS, 10μM mevastatin and 50μM mevalonate (chase). Coverslips were stained using Filipin and anti-CD63 antibody and analysed using a Zeiss LSM880 Confocal microscope with a 63x oil objective. Pearson’s correlation between Filipin and CD63 staining was determined using Volocity software (Perkin-Elmer) for 6 fields per sample with >8 cells per field. Statistical significance was determined using two-tailed Student’s T-Test with **p<0.01 and ***p<0.001 and error bars reflecting standard error of mean (SEM).

### Fluid phase endocytosis

Bulk endocytosis and delivery of extracellular materials to the late endosomal compartment was measured using the dye Sulforhodamine 101 (SR101, Sigma), following published protocols (42, 70). Briefly, Cells were seeded in live cell imaging dishes (MatTek), preincubated with Lysotracker Green (Molecular Probes) for 30 min, and SR101 dye was added directly to the cell medium at 25µg/ml. Samples were monitored by time-lapse on a Leica SP8 microscope adapted with a climate control chamber using a HCX PL 63x 1.32 oil objective and images taken at multiple positions at 2 min intervals. SR101 colocalisation with the Lysotracker-positive compartment is reported as Mander’s coefficient calculated using JACoP plug-in for ImageJ on the basis of 2 independent experiments. Total SR101 entering both cell lines was quantified at indicated time points by flow cytometry. Error bars indicate SD of the mean. Statistical evaluations report on Student’s T Test (analysis of two groups), with *p<0.05, **p<0.01, ***p<0.001.

### Electron microscopy

Samples were prepared for cryo sectioning as described elsewhere (71, 72). Briefly, C18orf8-deficient cells and wild-type cells were fixed for either 2h in freshly prepared 2% paraformaldehyde and 0.2% gluteraldehyde in 0.1M phosphate buffer, or for 24h in freshly prepared 2% paraformaldehyde in 0.1 M phosphate buffer. Fixed cells were scraped, embedded in 12% gelatin (type A, bloom 300, Sigma) and cut with a razor blade into 0.5 mm^3^ cubes. The sample blocks were infiltrated in phosphate buffer containing 2.3 M sucrose. Sucrose-infiltrated sample blocks were mounted on aluminium pins and plunged in liquid nitrogen. The vitrified samples were stored under liquid nitrogen.

Ultrathin cell sections of 75 nm were obtained essentially as described elsewhere (72). Briefly, the frozen sample was mounted in a cryo-ultramicrotome (Leica). The sample was trimmed to yield a squared block with a front face of about 300 x 250 μm (Diatome trimming tool). Using a diamond knife (Diatome) and antistatic devise (Leica) a ribbon of 75 nm thick sections was produced that was retrieved from the cryo-chamber with the lift-up hinge method (73). A droplet of 1.15 M sucrose was used for section retrieval. Obtained sections were transferred to a specimen grid previously coated with formvar and carbon.

Grids containing thawed cryo sections of cells fixed with 2% paraformaldehyde and 0.2% glutaraldehyde were incubated on the surface of 2% gelatin at 37 °C. Subsequently grids were rinsed to remove the gelatin and sucrose and were embedded in 1.8% methylcellulose and 0.6% uranyl acetate. In case of additional gold-labelling, sections of cells fixed with 2% paraformaldehyde were incubated on drops with 1 μM TNM-BODIPY for 30 minutes on ice, washed, blocked with 1% BSA in PBS, and then incubated with rabbit polyclonal anti-BODPIY antibody (Molecular Probes, Invitrogen, UK) (45). Sections were subsequently labelled with 10 nm protein A-coated gold particles (CMC, Utrecht University). EM imaging was performed with an Tecnai 20 transmission electron microscope (FEI) operated at 120 kV acceleration voltage.

### Identification of C18orf8 and NPC1 interaction partners by mass spectrometry (MS)

HeLa cells, or HeLa cells expressing 3xHA-C18orf8, C18orf8-3xHA or NPC1-3xHA were lysed in 1% digitonin, HA-tagged proteins were immune precipitated for 2.5 hours at 4°C using Ezview anti-HA agarose (Sigma-Aldrich) as described above, eluted using 0.5 μg/ml HA peptide (Sigma-Aldrich) and denatured using 2% SDS, 50mM Tris pH7.4. Eluted samples were reduced with 10mM TCEP for 10mins RT and alkylated with 40mM Iodoacetamide for 20mins RT in the dark. Reduced and alkylated samples were then submitted to digestion using the SP3 method (PMID: 25358341). Briefly, carboxylate coated paramagnetic beads are added to the sample and protein is bound to the beads by acidification with formic acid and addition of acetonitrile (ACN, final 50%). The beads are then washed sequentially with 100% ACN, 70% Ethanol (twice) and 100% ACN. 10uL of a buffer of TEAB (Triethylammonium bicarbonate) pH8 and 0.1% Sodium deoxycholate (SDC) is then added to the washed beads along with 100ng trypsin. Samples were then incubated overnight at 37 degrees with periodic shaking at 2000rpm. After digestion, peptides are immobilised on beads by addition of 200uL ACN and washed twice with 100uL ACN before eluting in 19uL 2% DMSO and removing the eluted peptide from the beads.

### MS data acquisition

Samples were acidified by addition of 1uL 10% TFA and the whole, 20uL sample injected. Data were acquired on an Orbitrap Fusion mass spectrometer (Thermo Scientific) coupled to an Ultimate 3000 RSLC nano UHPLC (Thermo Scientific). Samples were loaded at 10 μl/min for 5 min on to an Acclaim PepMap C18 cartridge trap column (300 um × 5 mm, 5 um particle size) in 0.1% TFA. After loading a linear gradient of 3–32% solvent B over 60 or 90min was used for sample separation with a column of the same stationary phase (75 µm × 75 cm, 2 µm particle size) before washing at 90% B and re-equilibration. Solvents were A: 0.1% FA and B:ACN/0.1% FA. MS settings were as follows. MS1: Quadrupole isolation, 120’000 resolution, 5e5 AGC target, 50 ms maximum injection time, ions injected for all parallelisable time. MS2: Quadrupole isolation at an isolation width of m/z 0.7, HCD fragmentation (NCE 34) with the ion trap scanning out in rapid mode from, 8e3 AGC target, 250 ms maximum injection time, ions accumulated for all parallelisable time. Target cycle time was 2s.

### MS data analysis

Spectra were searched by Mascot within Proteome Discoverer 2.1 in two rounds of searching. First search was against the UniProt Human reference proteome (26/09/17) and compendium of common contaminants (GPM). The second search took all unmatched spectra from the first search and searched against the human trEMBL database (Uniprot, 26/0917). The following search parameters were used. MS1 Tol: 10 ppm, MS2 Tol: 0.6 Da, Fixed mods: Carbamindomethyl (C) Var mods: Oxidation (M), Enzyme: Trypsin (/P). PSM FDR was calculated using Mascot percolator and was controlled at 0.01% for ‘high’ confidence PSMs and 0.05% for ‘medium’ confidence PSMs. Proteins were quantified using the Minora feature detector within Proteome Discoverer and values normalised against the median protein abundance of the whole sample.

## Supplementary figure legends

**Figure S1:**
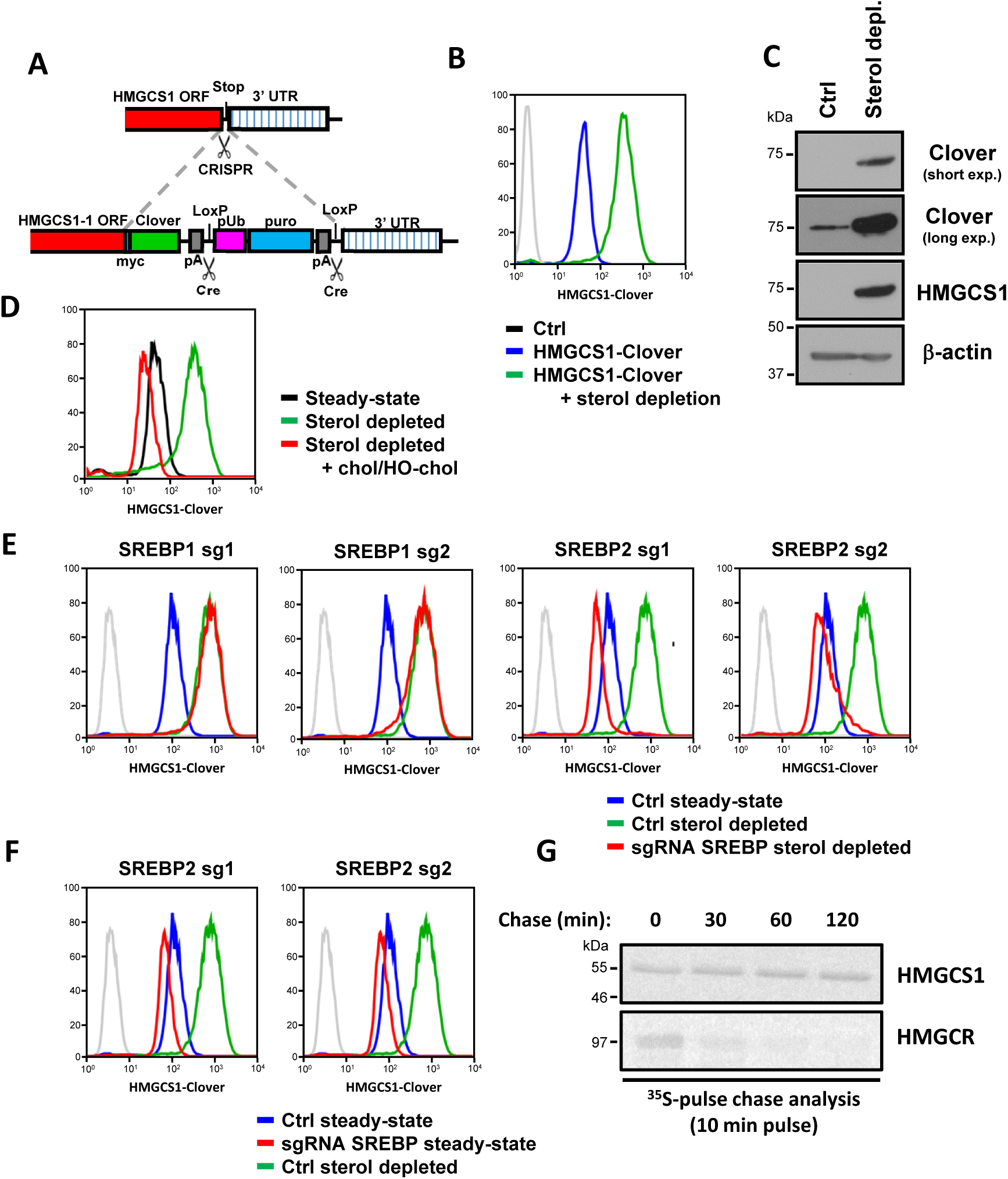
Characterisation of the HMGCS1-Clover cholesterol reporter (related to Fig 1). Creation of a HMGCS1-Clover CRISPR knock-in in HeLa cells. (**A**) Schematic for generation of an endogenous HMGCS1-Clover fusion protein. (**B**, **C**) Sterol depletion induces HMGCS1-Clover expression. HeLa HMGCS1-Clover cells were sterol depleted overnight and HMGCS1-Clover expression analysed by flow cytometry (**B**) or immune blotting using GFP (Clover) and HMGCS1-specfic antibodies. (**D**) Overnight supplementation with cholesterol and 25-hydroxy-cholesterol reverses HMGCS1-Clover upregulation in sterol-depleted HMGCS1-Clover cells (**E**, **F**) Sterol-depletion induced HMGCS1 up-regulation depends on SREBP2. HMGCS1-Clover CAS9 cells were transfected with two independent sgRNAs against *SREBF1*, *SREBF2* or both, sterol depleted overnight where indicated and analysed by flow cytometry for HMGCS1-Clover expression at day 8. (**G**) HMGCS1 protein stability is not cholesterol-sensitive. HeLa cells were sterol depleted overnight, pulse-labelled with ^35^S-Met/Cys and chased in the presence of excess cholesterol/25-hydroxy-cholesterol for the indicated times. After cell lysis, HMGCS1 or HMGCR was immune precipitated, separated by SDS-PAGE and analysed by autoradiography.

**Figure S2:**
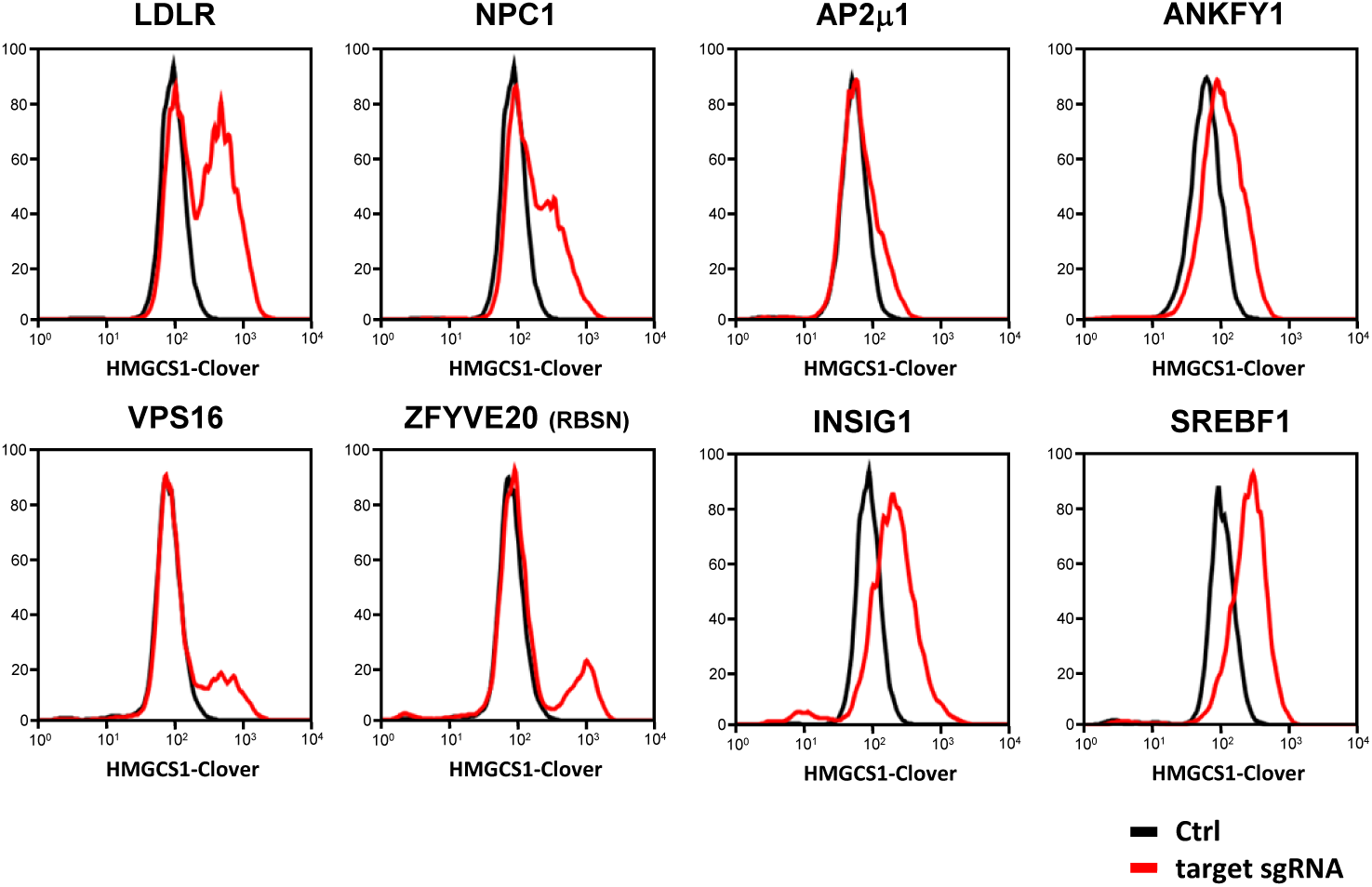
Validation of select genetic screening hits using the HMGCS1-Clover reporter (related to Fig 1). CRISPR knockout of select screening hits induces spontaneous HMGCS1-Clover up-regulation. HMGCS1-Clover CAS9 cells were transfected with sgRNAs against indicated targets and HMGCS1-Clover expression was analysed by flow cytometry at day 5 (*AP2μ1*), day 7 (*LDLR, NPC1, INSIG1*), day 8 (*ANKFY1*, *SREBF2*) or day 9 (*ZFYVE20, VPS16*).

**Figure S3:**
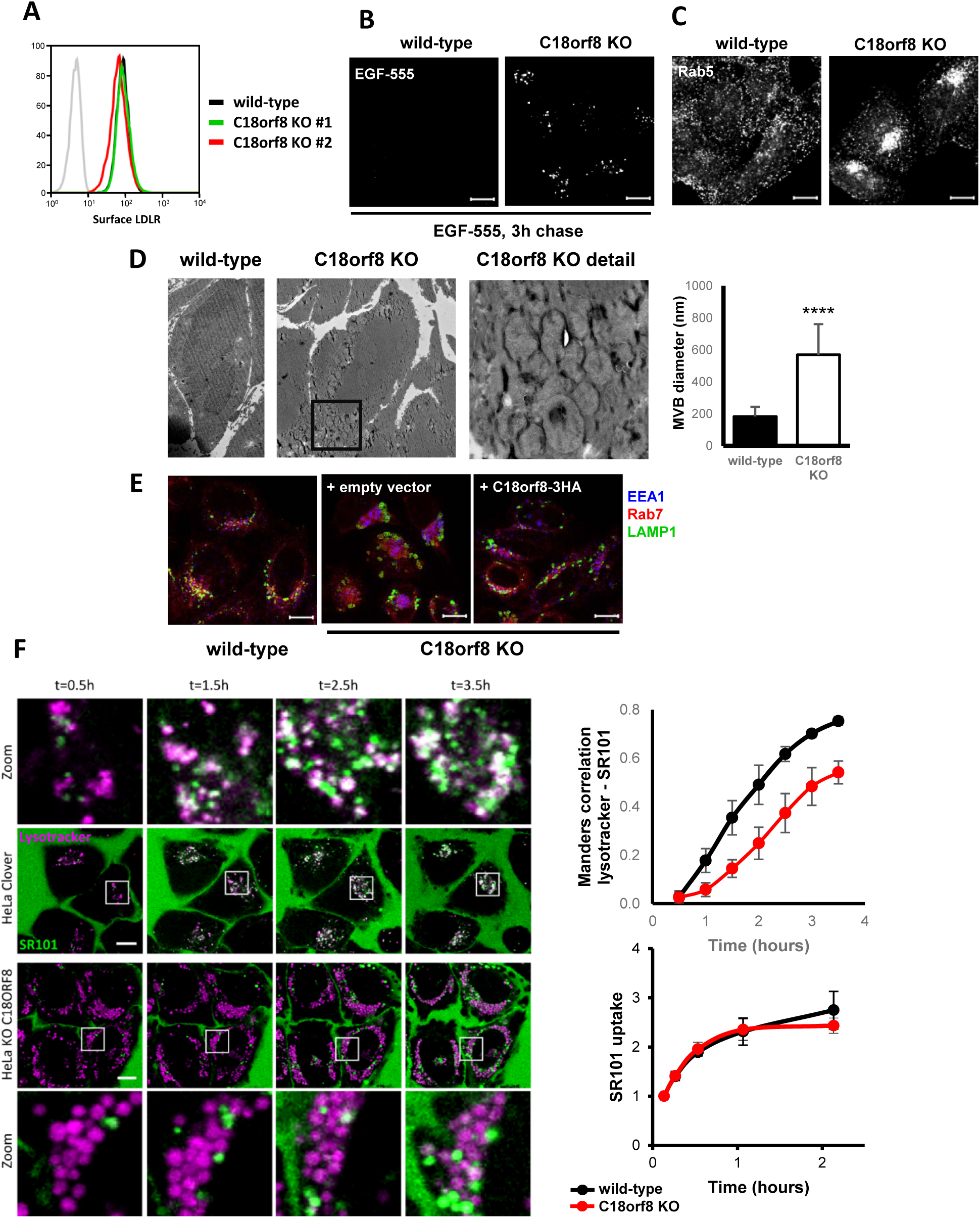
C*1*8orf8-deficient cells show severe defects in late endosome morphology and trafficking (related to Fig 2, 3). (**A**) *C18orf8*-deficient cells have normal cell surface LDL receptor (LDLR) expression. Wild-type or *C18orf8*-deficient cells were labelled for cell surface LDLR and analysed by flow cytometry. (**B**) *C18orf8*-deficient cells are defective in EGF degradation. Wild-type or *C18orf8*-deficient cells were pulse-labelled with AlexaFluor-555 conjugated EGF, incubated for 3 hours and fixed. Remaining EGF-555 fluorescence was visualised by confocal microscopy. (**C**) *C18orf8*-deficient cells show clustering of early endosomes (EE). Rab5 staining of wild-type and *C18orf8*-deficient cells. (**D**) C18orf8-deficient cells accumulated enlarged multivesicular bodies (MVBs). (MVBs). Electron microscopy of wild-type and *C18orf8*-deficient cells. MVB size was quantified in 30 cells (**** P<0.0001). (**E**) Complementation of endosomal phenotypes in *C18orf8*-deficient cells. Wild-type cells, C18orf8-deficient cells and C18orf8-deficient cells complemented with C18orf8-3xHA were labelled for EEA1, Rab7 and LAMP1 and visualised by confocal microscopy. (**F**) C18orf8-deficient cells show delayed SR101 trafficking into the late endosomal compartment. Wild-type or *C18orf8*-deficient cells were labelled with lysotracker (magenta), incubated with Sulforhodamine 101 dye (SR101, green) and imaged live for 3.5h at 1.5 min intervals. Select frame overlays from one of 5 positions imaged per experiment per condition are shown. Scale bars = 10µm. Quantification of SR101 trafficking into the lysotracker+ LE compartment over time is reported as Manders correlation (top right graph) determined from two independent experiment using 3-4 fields (containing 10-30 cells per field of view) per experiment (*** P<0.001). SR101 uptake was determined at indicated times by flow cytometry (bottom right graph).

**Figure S4:**
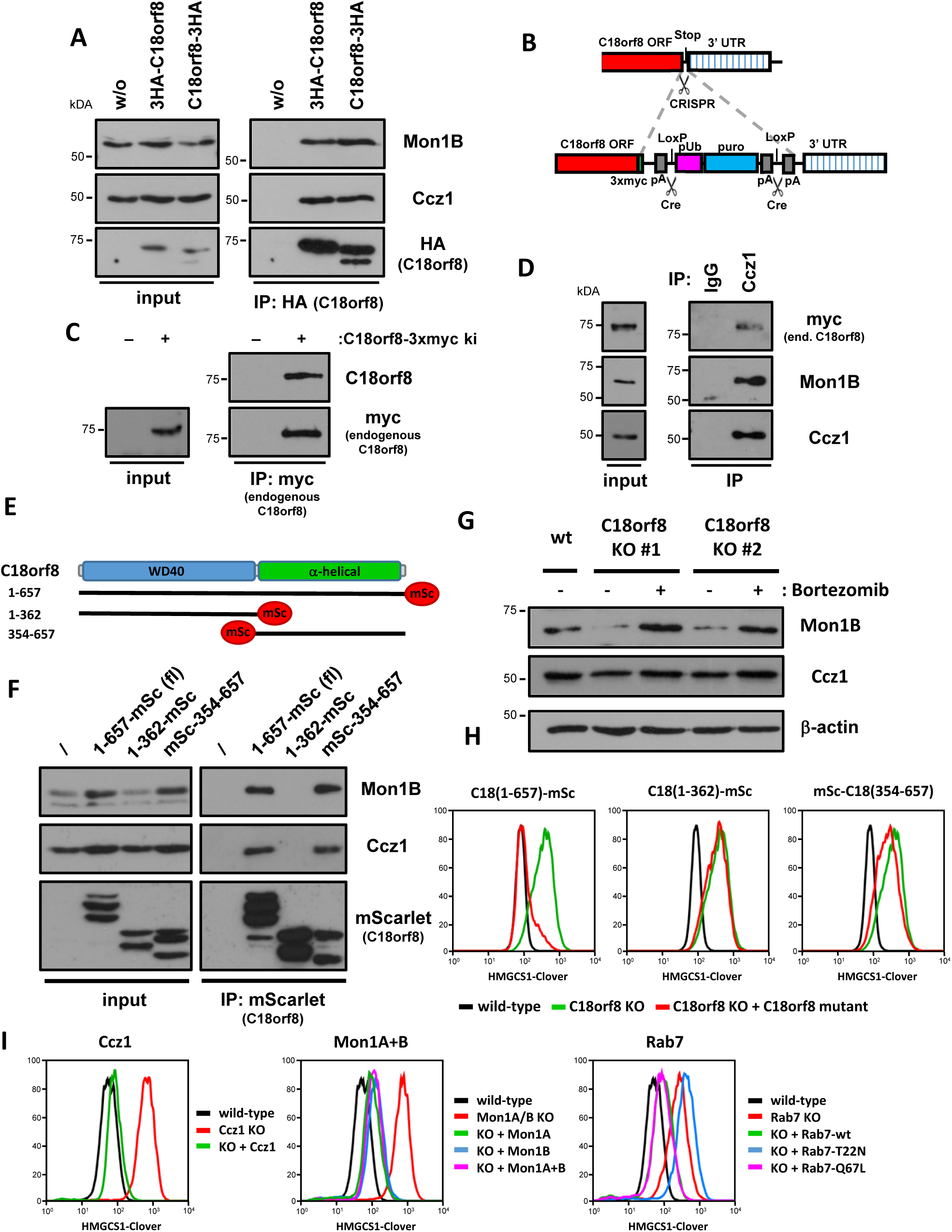
C18orf8 interacts with the mammalian Mon1-Ccz1 (MC1) complex and is essential for MC1 complex stability (related to Fig 4). (**A** – **C**) C18orf8 interacts with the mammalian MC1 complex. (**A**) Immune precipitation of overexpressed HA-tagged C18orf8 shows an interaction with Ccz1 and Mon1B. (**B**) Schematic for generation of an endogenous C18orf8-3xMyc fusion protein in HeLa cells. (**C**) Myc immune precipitation and immunoblotting of a knock-in clone shows the endogenous C18orf8-3xMyc fusion protein, detected by both C18orf8 and Myc antibodies. (**D**) Immune precipitation using a Ccz1-specific antibody but not an isotype control shows an interaction between Ccz1, endogenous C18orf8-3xMyc and Mon1B in C18orf8-3xMyc ki cells. (**E**, **F**) The C18orf8 C-terminal α-helical domain interacts with and stabilizes mammalian MC1. (**E**) Depiction of C18orf8 domain structure and mutants created. (**F**) C18orf8-deficient cells were complemented with mScarlet-labelled C18orf8 full-length (wt), C-terminal (AA 1-362) and N-terminal truncations (AA 354-657). mScarlet proteins were immune precipitated and immune precipitations and input controls were analysed by immune blotting using Mon1B, Ccz1 and mScarlet-specific antibodies. Note wild-type and N-terminal truncations (AA 354-657) stabilize Mon1B expression in input controls. (**G**) Mon1B expression in *C18orf8*-deficient cells is rescued by overnight proteasome inhibition (bortezomib). (**H**) The C18orf8 C-terminus alone is insufficient to complement *C18orf8* functions. *C18orf8*-deficient cells were complemented with mScarlet-labelled C18orf8 full-length (wt), C-terminal (AA 1-362) and N-terminal truncations (AA 354-657) and analysed by flow cytometry for HMGCS1-Clover expression. (**I**) Complementation of *Ccz1*-, *Mon1A/B*- and *Rab7*-deficient cells. *Ccz1*-, *Mon1A/B*- and *Rab7*-deficient cells with transduced with respectively Ccz1-3xMyc; Mon1A-3xMyc, Mon1B-3xMyc or both; and 2xHA-Rab7-wt, -T22N or –Q67L and HMGCS1-Clover expression was analysed using wild-type HMGCS1-Clover cells as a control.

**Figure S5:**
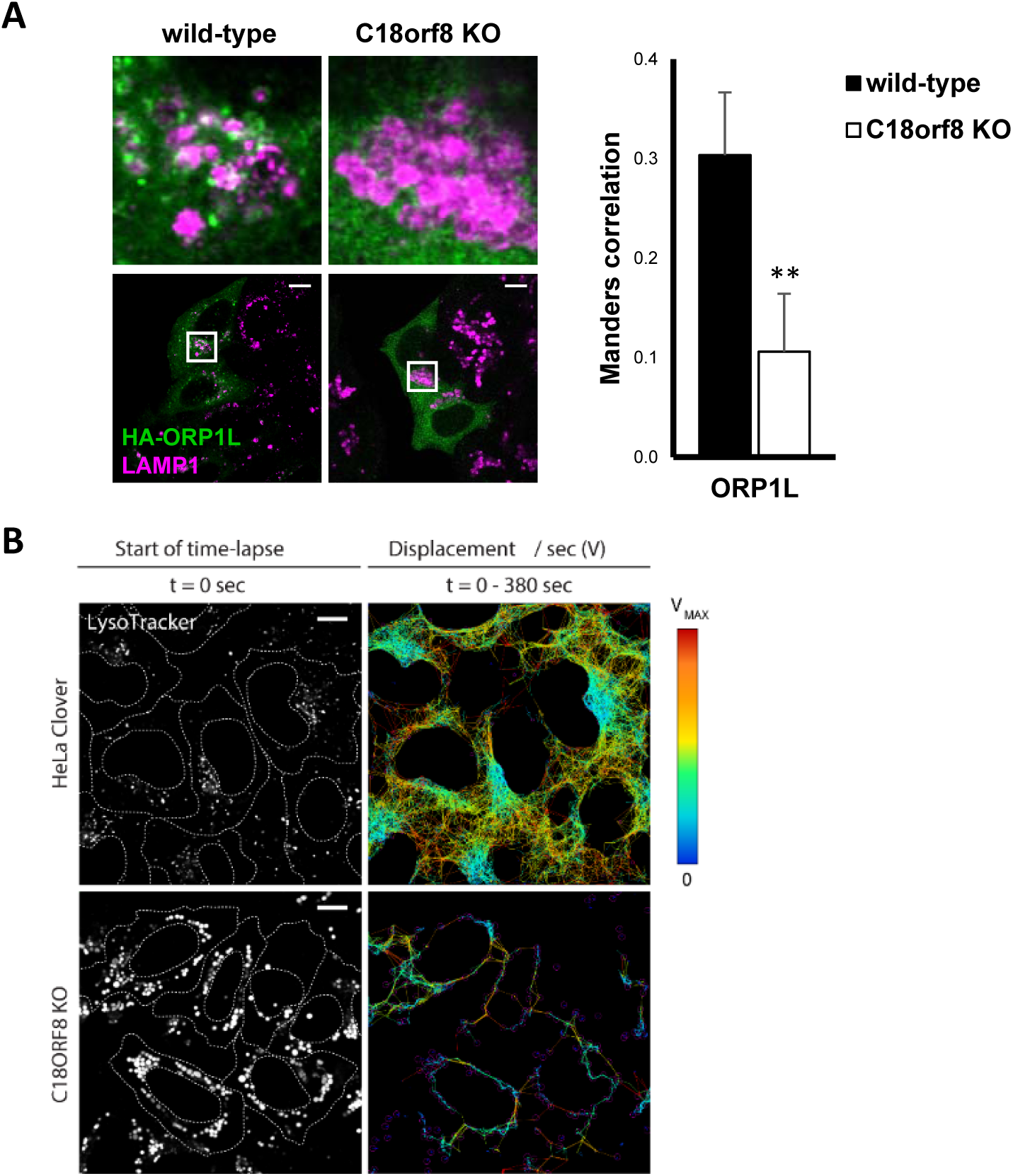
C*1*8orf8-deficient cells show impaired ORP1L recruitment and reduced late endosome motility (related to Fig 5). (**A**) Wild-type and *C18orf8*-deficient cells were transfected with HA-ORP1L and stained intracellularly for HA and LAMP1. Manders correlation was determined for 6-10 cells from two independent experiments (** P<0.01). (**B**) Wild-type or *C18orf8*-deficient cells were labelled with lysotracker and lysotracker+ endosomes (white) were traced for 380 sec using live cell microscopy at 3 sec intervals. Displacement of individual endosomes was calculated using TrackMate for Fiji and is depicted with maximum velocity of travel indicated by a heat map (blue: no motility, red: maximum motility attainable under selected tracking parameters).

**Figure S6:**
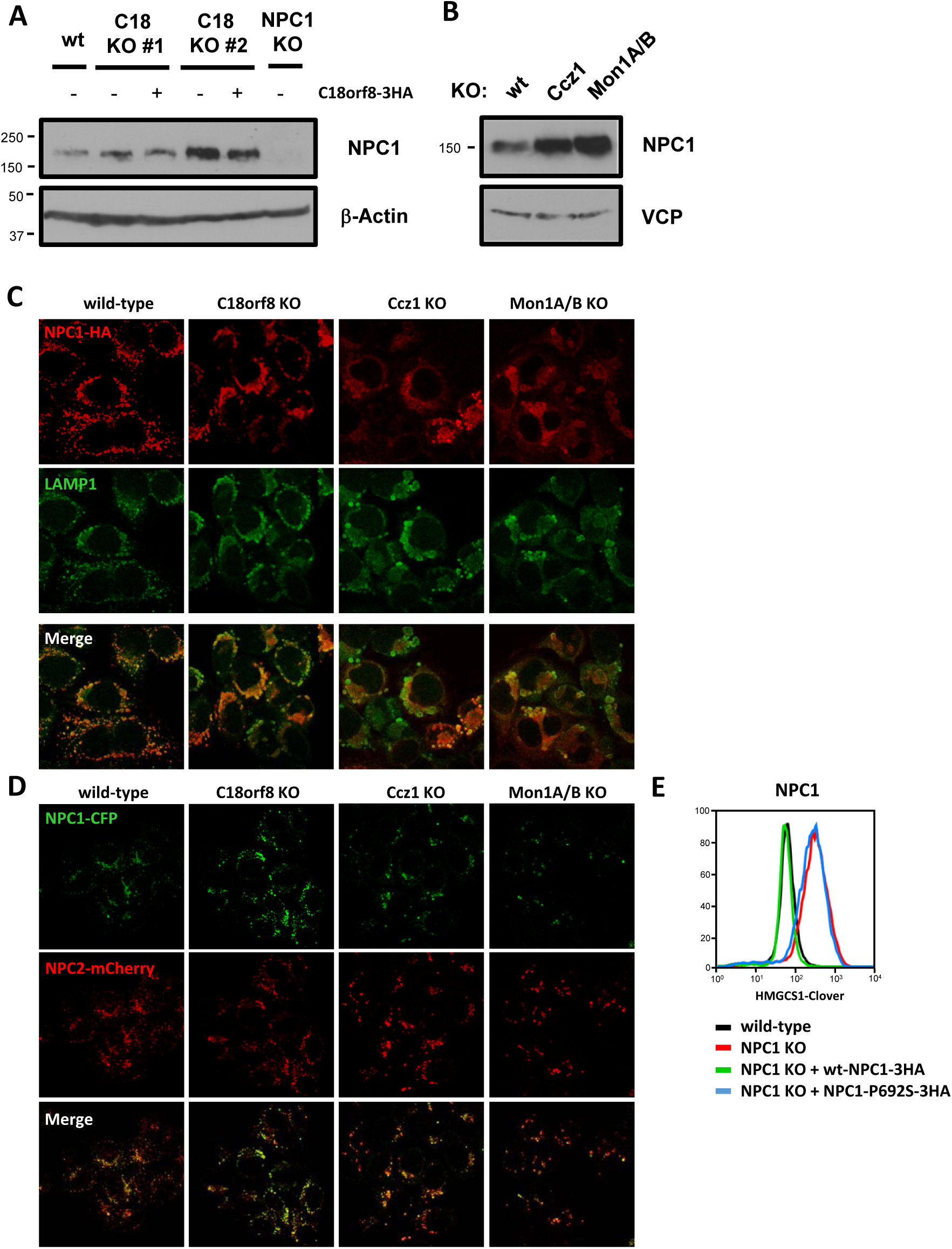
NPC1 expression and localisation is unperturbed in *C18orf8-*, *Ccz1-* and *Mon1A/B*-deficient cells (related to Fig 6). (**A, B**) *MCC*-deficient cells show normal to increased NPC1 expression. Immune blotting for endogenous NPC1 in (**A**) wild-type, *C18orf8*-deficient and complemented *C18orf8*-deficient cell clones; or (**B**) wild-type, *Ccz1*- and *Mon1A/B*-double deficient cells. (**C**, **D**) NPC1 localisation is largely unperturbed in *MCC*-deficient cells. (**C**) NPC1 localises to LAMP1+ LE/Ly in *MCC*-deficient cells. Wild-type, *C18orf8*-, *Ccz1*- and *Mon1A/B*-deficient cells were transduced with the inactive NPC1-P692S-3xHA and stained for HA and LAMP1. (**D**) NPC1 co-localises with the NPC2 cholesterol carrier in MCC-deficient cells. Wild-type, *C18orf8*-, *Ccz1*- and *Mon1A/B*-deficient cells were stably transduced with NPC1-P692S-CFP and NPC2-mCherry and monitored using confocal microscopy. (**E**) Complementation of NPC1-deficient cells. NPC1-deficient cells were transduced with wild-type or P692S mutant NPC1-3xHA and analysed by flow cytometry for HMGCS1-Clover expression.

**Figure S7:**
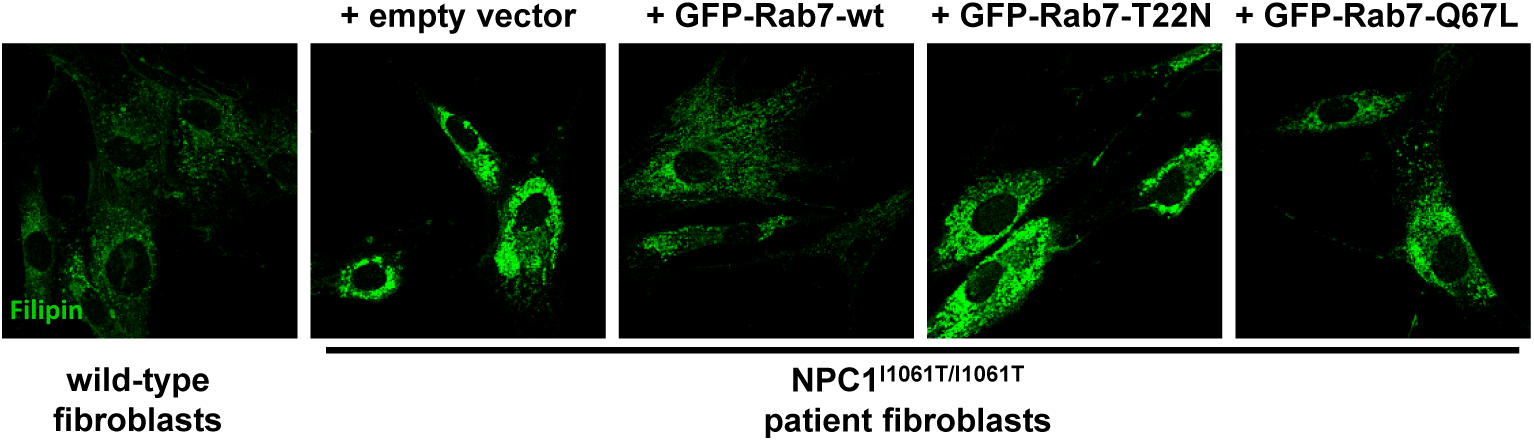
Rab7 overexpression reduces cholesterol accumulation in Niemann Pick primary patient fibroblasts (related to Fig 7). NPC1^I1061T/I1061T^ primary patient fibroblasts were stably transduced with GFP-Rab7 wild-type (wt), dominant negative (T22N) or constitutively active (Q67L) or empty vector and labelled for free cholesterol at day 7 using Filipin staining. Fibroblast from a healthy individual were used as a control. Overexpression of wild-type Rab7 reduces cholesterol accumulation in NPC patient fibroblasts, whereas Rab7-T22N does not and Rab7-Q67L gives an intermediate phenotype. Representative images from 2 independent experiments are shown.

**Table S1: Full RSA analysis of the HMGCS1-Clover screen for cholesterol regulators** (related to **Fig 1**). sgRNA enrichment for individual genes in the HMGCS1-Clover^high^ population compared to a library population, is indicated by RSA score and P-value. Genes enriched with p<10^-5^ are annotated into function pathways (See also Fig 1E, F).

